# Brain aging-dependent glioma traits reversible by NAD^+^/BDNF-mediated neuronal reactivation

**DOI:** 10.1101/2020.10.10.334748

**Authors:** Daisuke Yamashita, Victoria L Flanary, Rachel B Munk, Kazuhiro Sonomura, Saya Ozaki, Riki Kawaguchi, Satoshi Suehiro, Soniya Bastola, Marat S Pavlyukov, Shinobu Yamaguchi, Mayu A Nakano, Takeharu Kunieda, Dolores Hambardzumyan, Toru Kondo, Harley I Kornblum, David K Crossman, James R Hackney, Taka-aki Sato, Myriam Gorospe, Ichiro Nakano

## Abstract

The rise in aging population worldwide is increasing death from cancer, including glioblastoma. Here, we explore the impact of brain aging on glioma tumorigenesis. We find that glioblastoma in older patients and older mice displayed reduced neuronal signaling, including a decline of NTRK-like family member 6 (SLITRK6), a receptor for neurotrophic factor BDNF. This reduction was linked to the systemic decline of nicotinamide adenine dinucleotide (NAD^+^) with aging, as old mice exposed to young blood *via* parabiosis or supplemented with the NAD^+^ precursor NMN (nicotinamide mononucleotide) reverted phenotypically to young-brain responses to glioma, with reactivated neuronal signaling and reduced death from tumor burden. Interestingly, the phenotypic reversal by NMN was largely absent in old mice undergoing parabiosis with BDNF^+/-^ young mice and in BDNF^+/-^ mice undergoing tumor challenge, supporting the notion that the lower NAD^+^-BDNF signaling in the aging brain aggravated glioma tumorigenesis. We propose that the aging-associated decline in brain NAD^+^ worsens glioma outcomes at least in part by decreasing neuronal/synaptic activity and increasing neuroinflammation.

## Introduction

Aging is the biggest risk factor for glioblastoma, the most devastating form of the intra-parenchymal cancer glioma (*1–3*). Given the rapid global increase in the aging population, establishing more efficacious treatments for glioma patients is a social need and an area of great biomedical interest (*2–4*).

Glioblastoma cells were recently found capable of forming synaptic connections with neighboring neurons, thereby accelerating tumor growth (*5–7*). This striking discovery highlights a mechanism operating in young gliomagenesis, given that the data were largely collected in young mouse tumor models. In aging brains, on the other hand, marked decreases of neuronal number and activity trigger initiate and promote many diseases, often exacerbated by chronic inflammation (*8, 9*). Importantly, the rise in chronic inflammation is more apparent in the cerebral cortex and white matter, frequent locations where glioma arises in adults, and may play a causal role in glioma as it does in a variety of other human cancers (*10, 11*). Of note, resistance to therapy in gliomas is further accelerated by local chronic inflammation, which creates a harsh tumor microenvironment following surgery, chemotherapy, or radiotherapy, and worsens the prognosis for patients. Despite many advances in the knowledge of glioma development and therapy, our understanding of aging-related molecular and phenotypic changes remains shallow, preventing the implementation of practice-changing discoveries in the clinic, particularly for older glioblastoma patients.

The aging-associated decline in the activity of nicotinamide adenine dinucleotide (NAD^+^) has been implicated in pathologies including cancer (*12, 13*). NAD^+^ is an essential metabolite in the brain, primarily involved in energy homeostasis and DNA repair (*13*). Elevating NAD^+^ levels through administration of its precursor nicotinamide mononucleotide (NMN) had beneficial effects against aging-related physiological declines and diseases(*14*), and boosting NAD^+^ levels extended the lifespan of laboratory animals including rodents(*14, 15*). However, whether NAD^+^ activation leads to the prevention or therapeutic improvements in cancer has not been reported.

The glioblastoma-brain parenchyma interface (termed ‘tumor edge’) creates a unique ecosystem where several somatic cells (e.g. vascular endothelial cells and astrocytes) contribute to elevating tumor malignancy. In particular, an astrocyte-derived decline in NAD^+^ activity appears to affect glioblastoma aggressiveness at the tumor edge, at least in young mouse models. Nonetheless, the effect of NAD^+^ activity on tumorigenesis in the ecosystem of older brains has not been elucidated yet.

In this study, we addressed the hypothesis that rejuvenating brains by supplementation of certain circulating factors including NMN might reverse the glioblastoma-permissive trait of aging brains. Using mice bearing glioblastoma, we present evidence that treatment with NMN or exposure to circulating factors from young mice reactivated neuronal signaling and reduced tumor-causing death. We identify BDNF as an effector of these beneficial interventions and propose that the rise of aggressive glioblastoma with age is linked to a concomitant decline in brain NAD^+^ that reduces neuronal function and increases neuroinflammation.

## Results

### Brain aging exacerbates glioma aggressiveness

Comparison of the distribution of high-grade gliomas (HGG) [WHO Grade III and Grade IV (glioblastoma)] between younger (<60) and older (>60) patients using the Cancer Genome Atlas (TCGA) database revealed an elevation of glioblastoma in the older population; Grade III astrocytomas were more prevalent in the young population (**Fig. 1A**). Meta-analysis across four different datasets (TCGA, Rembrandt, GSE36245, and GSE13041) showed significantly reduced overall survival (OS) and progression-free survival (PFS) in elderly HGG patients compared to younger patients (*16, 17*) (**Fig. 1B** and fig. S1, A-F). Of note, the reduced overall survival of older patients was not due to IDH (Isocitrate dehydrogenase) status, as a significantly poorer prognosis was seen for older patients in both IDH wild-type (WT) and mutant groups within the TCGA dataset (**fig. S1B**). In addition, the relative distribution of subtypes of IDH WT tumors in the TCGA database [proneural (PN), mesenchymal (MES), and classical (CL)] was the same in both age groups (**fig. S1G**). Gene ontology (GO) analysis of the RNA-seq data indicated that younger tumors expressed mRNAs encoding proteins implicated in neuronal and synaptic pathways (e.g. neuropeptide receptor activity and neuron projection), whereas older tumors expressed mRNAs encoding proteins implicated in neuronal inflammation and DNA methylation (e.g. oxidative stress-induced senescence and DNA damage/telomere stress-induced senescence) (**Fig. 1, C and** D**; fig. S2, A and B**).

**Figure 1.**
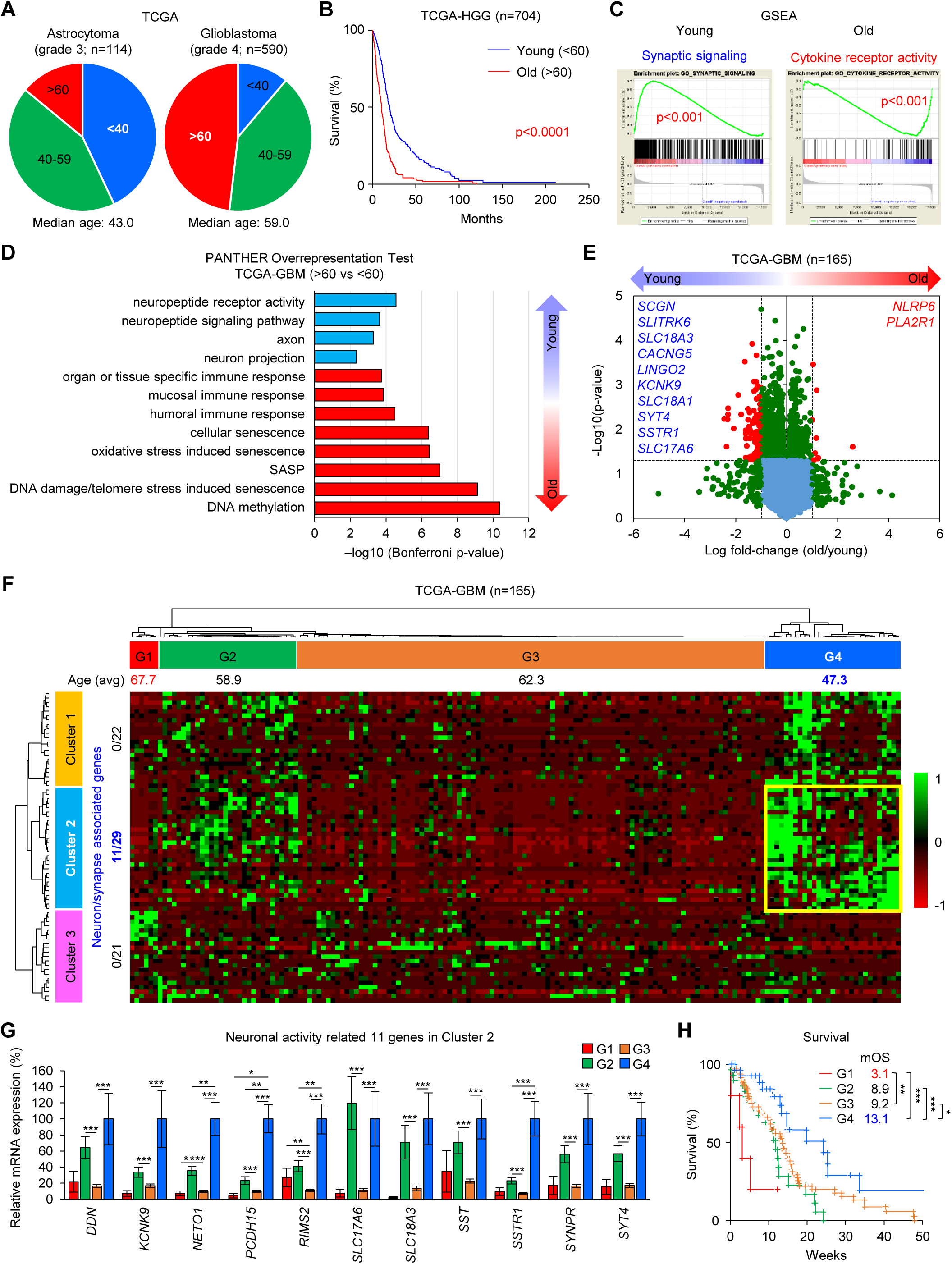
Brain aging enhances the aggressiveness of glioblastoma tumors. (**A**) Pie chart comparing median ages of patients with WHO Grade III astrocytoma (43.0) and glioblastoma (59.0) according to the TCGA dataset. n=114 and 590, respectively. (**B**) Kaplan-Meier analysis comparing overall survival of young and old glioblastoma patients in the TCGA-IDH-wild-type dataset (n=368). *p*<0.0001. (**C**) Gene set enrichment analysis (GSEA), highlighting synaptic signaling and cytokine receptor activity in young and old glioblastoma patients, respectively. Gene expression profiles were obtained from the TCGA RNA-seq database. (**D**) Bar graph comparing the pathways activated in young and old glioblastoma patients in the TCGA dataset. (**E**) Volcano plot displaying the most upregulated genes in young and old glioblastomas in the TCGA dataset. (**F**) Unbiased heat map clustering of RNA-seq analysis, TCGA dataset. (**G**) RNA-seq analysis of the 11 neuron/synapse-associated genes in cluster 2 in (**F**). (**H**) Kaplan-Meier analysis comparing the 4 patient groups in (**F**) (n=165; 5 G1, 28 G2, 85 G3, and 27 G4).

The genes upregulated in younger tumors were largely related to neuronal/synaptic activity (i.e. *SCGN*, *SLITRK6*, and *SLC18A3*) (**Fig. 1E**). In addition, the unbiased clustering analysis of the RNA-seq data identified 4 previously unidentified patient groups (G1-4) (**Fig. 1F**). Of note, none of the clusters correlated with any of the subtypes – PN, MES, or CL(*18*). instead, there was clustering with age: the youngest glioblastoma patients (Group 4, G4) (average age 47.3 y.o.) harbored distinct gene expression patterns with 11 markedly elevated genes implicated in neuronal activity that accumulated in Cluster 2, while Group 1 (G1) comprised the oldest glioblastoma patients (67.7 y.o.) and its gene expression pattern was substantially different and almost mutually exclusive from that of G4 (**Fig. 1F and** G). Importantly, the overall survival of G4 was significantly better than the others, and the worst overall survival was seen for G1 (**Fig. 1H, and fig. S3A-C**). Among the 11 genes representative of G4, seven displayed individual survival benefits in the Rembrandt database for WHO Grade II to IV gliomas, indicating that neuron- and synapse-related genes were likely associated with lower grades of glioma and better patient outcomes (**fig. S4, A and B**). Collectively, these findings suggest that transcriptomic programs toward a less neuronal and more neuroinflammatory phenotype contribute to the aging-related increases in glioma grade, phenotypic aggressiveness, and patient mortality. We also suggest that a new transcriptomic subclassification termed “Neuronal Activity-Based (NAB)” may be more clinically relevant than the currently used transcriptional subtyping, given the direct link to the patient’s post-surgical life expectancy.

### Mouse brain tumor models recapitulate the aging-associated phenotypic shift in human glioblastoma

Given these clinical findings, we next investigated mouse tumor models for their aging-related changes. We first transplanted mouse glioblastoma cells into young (2 months old (m.o.)) and old (16 m.o.) mice. Specifically, we used 4 mouse glioblastoma models. The first three, the glioma sphere lines ms7080, AR006, and ms6835, were established from glioblastoma-like tumors forming spontaneously in mice by ablation of genes *Tp53*, *Pten*, and *Nf1* (ms7080)(*19*), *CreER; Pten*^loxP/loxP^; *Tp53*^loxP/loxP^; *Rb1*^loxP/loxP^ (*20*), and *Cre; Nf1*^f/f^ and *p53*^f/^(*21*), respectively. The fourth, NSCL61, was established from p53-deficient murine neural stem cells overexpressing oncogenic HRas^L61^(*22*). Following intracranial injection and tumor establishment with all four models in young and old mice, we stained the tumor-bearing mouse brains with hematoxylin and eosin (H&E) to assess their histopathology. All these mouse tumors displayed pseudo-palisading adjacent to central necrosis and microvascular proliferation, two histopathological hallmarks of human glioblastoma (**fig. S5A**). As shown by H&E staining of the representative m7080-derived tumors, old mice harbored densely packed tumor cells with large central necrosis, while young mouse tumors exhibited relatively lower cellularity and noticeably smaller necrotic areas (**Fig. 2A**). The survival time of older tumor-bearing mice was much shorter than that of younger mice in all four models following tumor challenge (**Fig. 2, B and C**). Additionally, comparison of the xenograft models utilizing the patient-derived glioma spheres (1051x1) injected into young *vs.* old immunocompromised SCID mice yielded similar results (**Fig. 2C** and **fig. S5C**).

**Figure 2.**
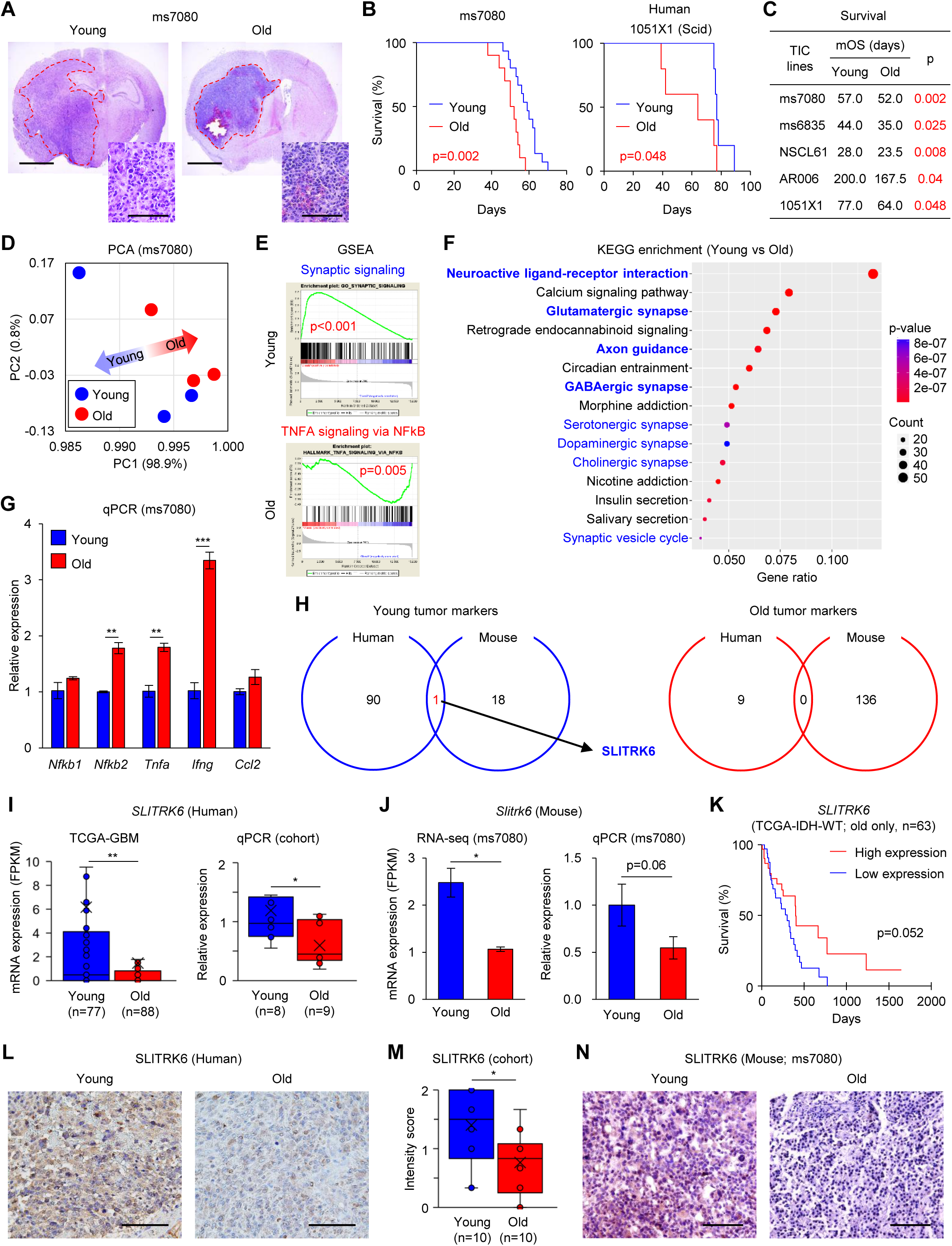
Tumors in aged mouse brains recapitulate human glioblastoma phenotypes. (**A**) Hematoxylin and eosin (H&E) staining of young (left) and old (right) mouse brains following intracranial injection of ms7080 cells. Scale bars represent 2 mm (full brain section) and 100 µm (enlarged section). (**B**) Kaplan-Meier analysis comparing young and old mice following intracranial injection of ms7080 cells (left; n=15 for young; n=10 for old) and young and old Scid mice following intracranial injection of human 1051X1 cells (right; n=5 for young and old). (**C**) Comparison of the median survival days of young and old mice following intracranial injection in 4 mouse glioblastoma models (ms7080-AR006) and 1 patient-derived tumor model (1051X1). (**D**) Principal component analysis (PCA) of RNA-seq data comparing young and old tumors derived from ms7080 cells. (**E**) GSEA displaying the indicated pathways activated in young and old ms7080 tumors. (**F**) KEGG enrichment analysis of RNA-seq data for comparison between young tumors and old tumors. (**G**) RT-qPCR analysis of inflammatory markers in young and old ms7080 tumors. Data are the means ± SD (n=3). \*\**p*<0.01, \*\*\**p*<0.001. (**H**) Venn diagram showing shared upregulated mRNAs in young and old glioblastomas in TCGA data and ms7080 tumors. (**I**) *SLITRK6* mRNA expression levels in human glioblastoma tissues, obtained from the TCGA dataset (left) and by RT-qPCR data (right; n=8 for young; n=9 for old). \*\**p*<0.01. (**J**) RNA-seq (left) and RT-qPCR (right) analyses of *Slitrk6* mRNA levels in young and old ms7080 tumors (n=9 for young and old). \**p*<0.05. (**K**) Kaplan-Meier analysis of OS of old glioblastoma patients in the TCGA dataset (n=63). (**L**) Representative IHC analysis of SLITRK6 signals in young and old human glioblastoma tissues. Scale bar represents 100 µm. (**M**) Intensity scoring of SLITRK6 signals detected by IHC in young and old human glioblastoma tissues (n=10 for young and old). (**N**) Representative IHC images for SLITRK6 in young and old mouse tumor tissues. Scale bar represents 100 µm.

To identify the molecular mechanisms underlying the aging-associated phenotypic shift in tumors, we performed RNA-seq analysis of these mouse tumors. Principal component analysis (PCA) harboring PC1 as the identity determinant for 98.9% demonstrated an axis of gene expression for tumors from young to old mice (**Fig. 2D**). Similar to the data with glioblastoma patients (Fig. 1, C **and** D), GSEA identified an elevated activity of synapse organization pathways in younger tumors, while older tumors displayed elevations in cytokine production and inflammatory response pathways (**Fig. 2E** and **fig. S6A**). KEGG analysis confirmed that neuronal/synaptic pathways were more activated in younger tumors (**Fig. 2F** and **fig. S6, B and C**) and RT-qPCR analysis confirmed the elevation of inflammatory markers in older tumors (e.g. *Ifng*, *Tnfa*, and *Nfkb2* mRNAs) (**Fig. 2G**).

We then compared the gene expression profiles in young and old mouse tumors with those in human glioblastomas in the TCGA database. As shown in **Fig. 2H**, *SLITRK6* mRNA, encoding a membrane-bound protein that is known to promote proliferation of peripheral neurons within the inner ear and eye (*23*), was the only transcript significantly elevated in both humans and mice. RT-qPCR analysis of our cohort of 17 glioblastoma patients confirmed that *SLITRK6* mRNA expression was in fact significantly higher in younger tumors (**Fig. 2I**). RNA-seq and RT-qPCR analyses of mouse tumors showed similar results (**Fig. 2J**). As expected, the TCGA database revealed shorter overall survival of *Slitrk6* low-expression group in older glioblastoma patients (**Fig. 2K**). Immunohistochemistry analysis of human glioblastomas and mouse tumors revealed higher protein expression levels for SLITRK6 in younger tumors (**Fig. 2, L to N**). Taken together, these data suggest that the mouse tumor models recapitulate the age-associated phenotypic and molecular reprogramming seen in human glioblastoma, with SLITRK6 as the only shared molecule declining with aging in mouse and patient tumors.

### Exposure of old brains to young blood alters the brain microenvironment, diminishing glioblastoma aggressiveness

We then asked if the aging-associated difference in tumor aggressiveness was due to the presence of different circulating factors in young *vs.* old mice affecting the architecture of brain microenvironment. To address this question, we performed a series of parabiosis experiments, physically connecting a young mouse (2 m.o.) and an old mouse (16 m.o.) to share blood for 5 weeks; following disconnection, these mice received tumor challenges (**Fig. 3A**). In control experiments, we paired two young control mice (termed ‘Young-cont’) and old two mice (termed ‘Old-cont’). While both Young-cont and Old-cont developed almost identical tumors to those without parabiosis, old mice receiving young blood (termed ‘Old-para’) exhibited substantial changes in tumor morphology and cellular architecture, resembling those of Young-cont (**Fig. 3B** and fig. S7, A and B). Accordingly, we also observed a remarkable improvement in survival of Old-para in all three glioblastoma models (**Fig. 3C**). Surprisingly, young mice receiving old blood (termed ‘Young-para’) developed tumors that appeared strikingly similar to those found in Old-cont (**fig. S7C**). At one month following injection of ms7080 cells, H&E staining of Old-para showed relatively less dense tumor cells with infiltration to the corpus callosum and lateral ventricle, which resembled those in Young-cont (**fig. S7D**). Consistent with the data with human glioblastoma patients and mouse models without parabiosis (**Fig. 2F**), KEGG analysis of RNA-seq data of Old-para revealed substantially overlapping elevation of neuronal/synaptic pathways (**Fig. 3D** and **fig. S8, A and B**). In addition, *Slitrk6* mRNA levels were markedly increased in Old-para, as determined by both RNA-seq and RT-qPCR analyses (**Fig. 3, E and F**). Furthermore, IHC analysis of Old-para displayed higher SLITRK6 signals comparable to those in Young-cont (**Fig. 3G**).

**Figure 3.**
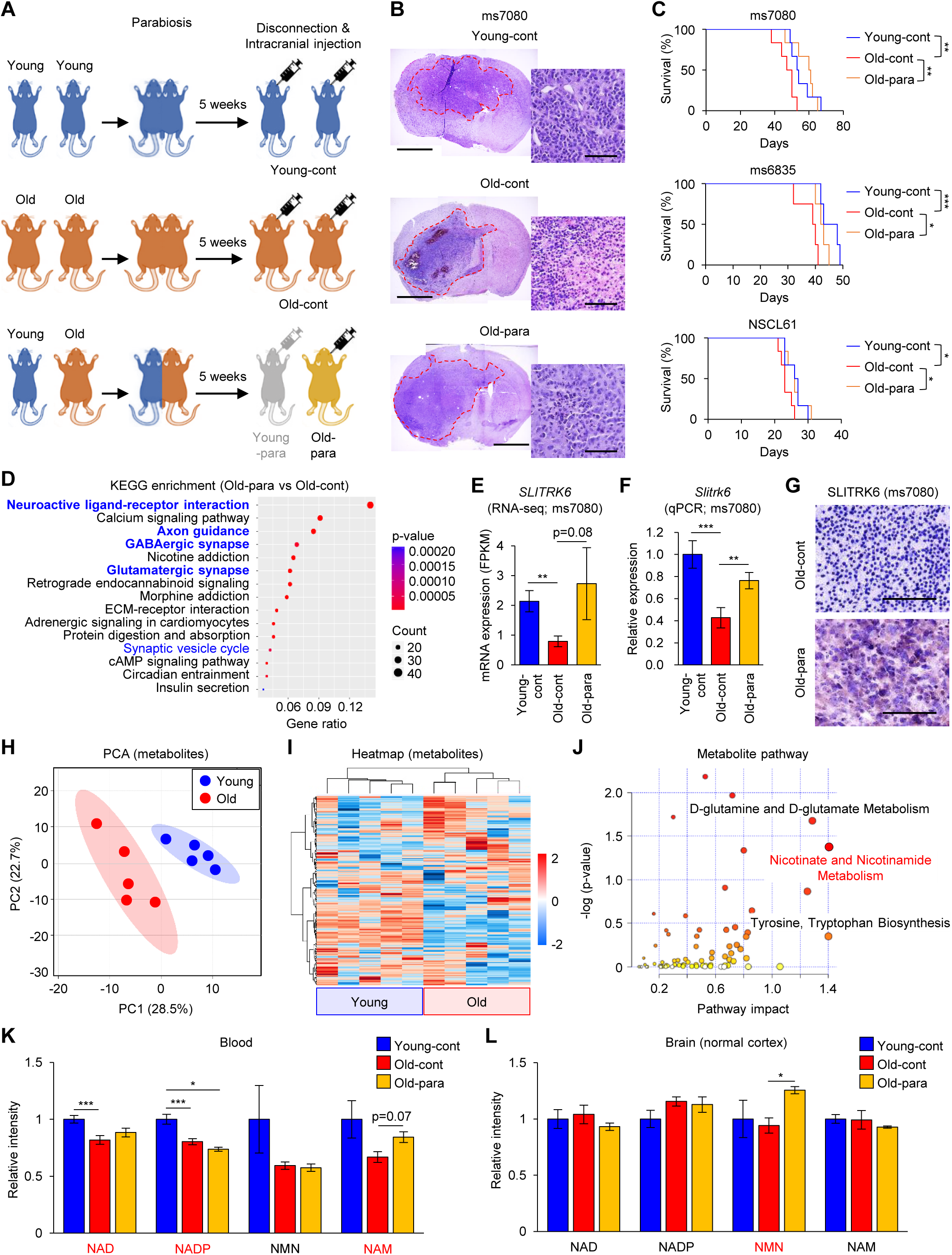
Pre-exposure of old brains to young blood reverses glioblastoma phenotype. (**A**) Schematic of the chronological order for parabiosis connecting young and old mice, followed by disconnection and tumor challenge (ms7080). (**B**) H&E staining of the brains indicated. Scale bars represent 2 mm (left) and 100 µm (right). (**C**) Kaplan-Meier analysis comparing the survival of each mouse group following injection of ms7080, ms6835, and NSCL61 (n=18, 12, 12 total, respectively). \**p*<0.05, \*\**p*<0.01, \*\*\**p*<0.001. (**D**) KEGG enrichment analysis of RNA-seq data for comparison between Old-para and Old-cont tumors. (**E-F**) *Slitrk6* mRNA expression levels by RNA-seq (**E**) and RT-qPCR (**F**) analyses of mouse tumors. Data are the means ± SD (n=3 each). \*\**p*<0.01, \*\*\**p*<0.001. (**G**) Representative IHC analysis of SLITRK6 expression levels in old-cont and old-para mouse tumors. Scale bars represent 100 µm. (**H**) PCA comparing blood samples from young and old mice. (**I**) Heat map of LC/MS data with indicated blood samples (n=5 for young and old). (**J**) Metabolite pathways upregulated in young blood. (**K**) Quantitative analysis of NAD, NADP, NMN, and NAM levels in blood, measured by LC/MS (n=14 for young-cont and old-cont; n=3 for old-para). \**p*<0.05, \*\*\**p*<0.001. (**L**) Quantitative analysis of NAD^+^, NADP, NMN, and NAM levels in normal cerebral cortices, measured by LC/MS (n=9 for young-cont; n=8 for old-cont; n=3 for old-para). \**p*<0.05.

Next, in order to determine the circulating factors responsible for these phenotypic shifts, we performed mass spectroscopy analysis of metabolites in blood obtained from young and old mice. PCA demonstrated an axis of metabolite expression from young to old samples (**Fig. 3H**). Also, Heat map clustering analysis showed a change in overall metabolite expression profiles between young and old samples (**Fig. 3I**). Metabolite pathway analysis identified the nicotinate and nicotinamide metabolism pathway as one of the most activated pathways in young blood (**Fig. 3J**). Based on these results, we measured the expression of NAD^+^-related metabolites in blood and normal cortical tissues obtained from young and old mice, as well as post-parabiosis mice. This analysis revealed that NAD^+^ and nicotinamide adenine dinucleotide phosphate (NADP) levels were among the most upregulated in Young-cont blood samples as compared to Old-cont, while Old-para restored some, if not all, nicotinamide (NAM) levels (**Fig. 3K**). Consistent with these data, nicotinamide mononucleotide (NMN) was significantly increased in post-parabiotic normal cortical tissues (**Fig. 3L**). Collectively, these findings suggest that young blood contains circulating factors including those in the NAD^+^ pathway (including NMN, which penetrates the blood-brain barrier) that might be capable of reversing the aging-associated tumor aggressiveness and the loss of neuronal activation in the brain tumors.

### Systemic treatment with NMN diminishes tumor aggressiveness

To test this possibility, we investigated whether direct activation of the NAD^+^ pathway in older brains was capable of reversing the aging-dependent aggravation of glioblastoma. To activate the NAD^+^ pathway, NMN was systemically administered to old WT mice (16 m.o.) for 3 weeks. Treated mice underwent tumor challenge with 4 murine and 1 human glioblastoma models (**Fig. 4A**). As shown, NMN pre-treatment of old mice (termed ‘NMN-before-tumor’) led to fewer pseudo-palisading or microvascular proliferation areas in tumors determined by H&E staining, indicating a shift of the subsequent tumor grade from glioblastoma to Grade III gliomas (**Fig. 4, B and C and fig. S9, A and B**). To examine this possibility further, double-blinded assessments of tumor grading was performed by a board-certified neuropathologist (JRH), who independently determined that a higher number of old mouse tumors were congruent with a diagnosis of glioblastoma, unlike the tumors in young mice and those in NMN-before-tumor mice (**fig. S9C**). Consistent with this observation, NMN pre-treatment conferred significantly longer survival in all 4 tested models (**Fig. 4D** and **fig. S9D**). In sharp contrast to these changes, NMN post-treatment (termed ‘NMN-after-tumor’) did not confer any survival benefits or histopathological changes (**Fig. 4E** and **fig. S9E**).

**Figure 4.**
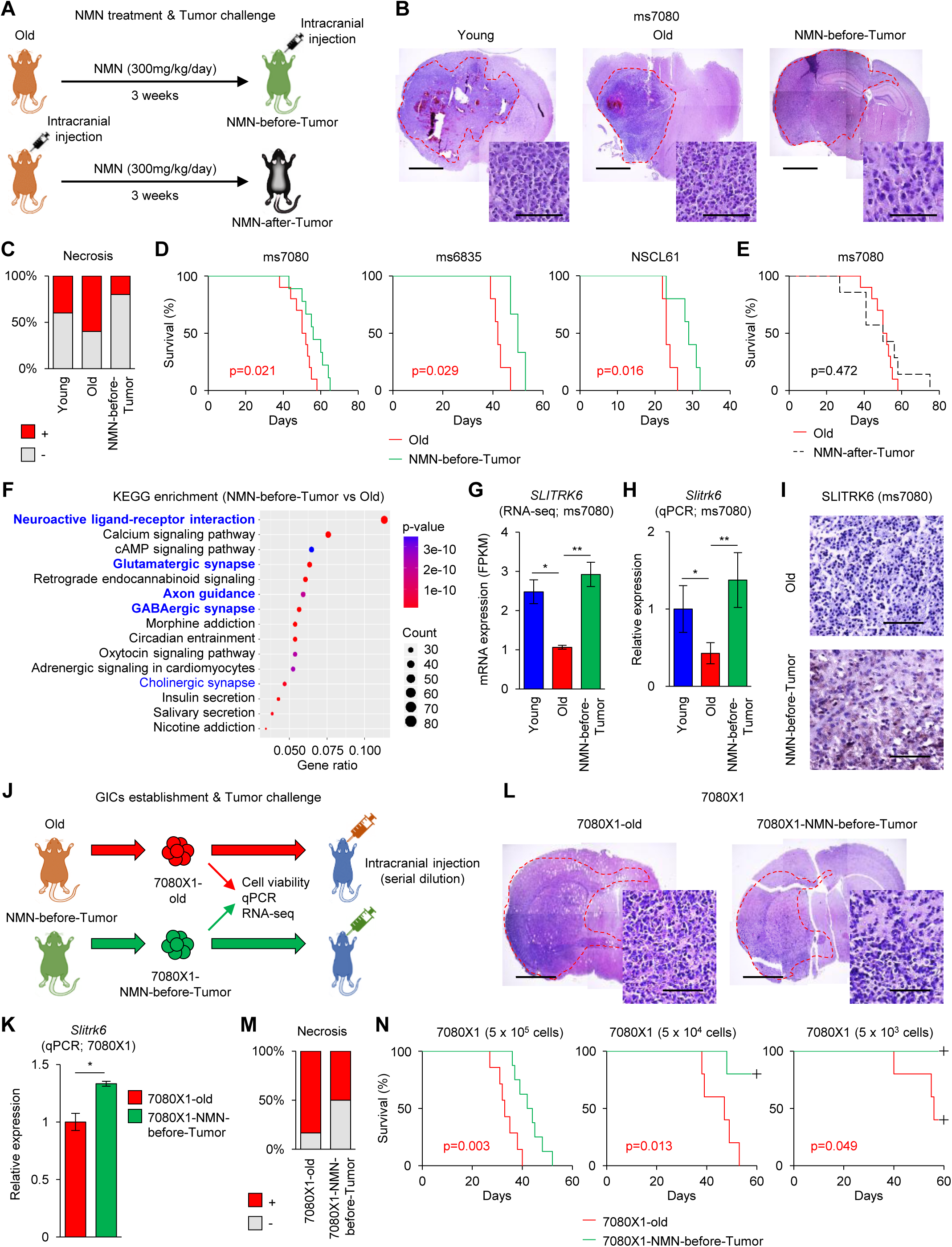
Systemic pre-treatment with NMN diminishes tumor aggressiveness. (**A**) Schematic of the timing of NMN administration before tumor challenge (ms7080) in 16 m.o. mice. (**B-D**) Representative H&E staining (**B**), rate of necrosis (**C**), and Kaplan-Meier survival analysis (**D**) of the indicated mouse brains. In (B) scale bars represent 2 mm (whole brain) and 100 µm (enlarged field); in (C) n=5 mice per group; in (D) *left*, n=10 for old, n=9 for NMN-before-tumor with ms7080 tumors; *middle*, n=5 for old, n=5 for NMN-before-tumor with ms6835 tumors; and *right*, n=5 for old and NMN-before-tumor with NSCL61 tumors. (**E**) Kaplan-Meier survival analysis of mice with ms7080 tumors treated with NMN after tumor challenge (n=10 for old; n=7 for NMN-after-tumor). (**F**) KEGG enrichment analysis of RNA-seq data for comparison between old tumors and NMN-before-tumor tumors. (**G-H**) *Slitrk6* mRNA levels by RNA-seq (G) and RT-qPCR (H) analyses of ms7080 tumor. Data are the means ± SD (n=3). \**p*<0.05, \*\**p*<0.01. (**I**) Representative IHC images for SLITRK6 in ms7080 tumors (old and NMN-before-tumor). Scale bars represent 100 µm. (**J**) Schematic representation of the experiments in (**K**-**N**). (**K**) RT-qPCR analysis of the levels of *Slitrk6* mRNA in 7080X1-old and -NMN-before-tumor cells. Data are the means ± SD (n=3). \**p*<0.05. (**L**) Representative H&E staining of mouse brains following intracranial injection of 7080X1-old and -NMN-before-tumor cells. Scale bars represent 2 mm (left) and 100 µm (right). (**M**) Rate of necrosis in 7080X1 tumors (7080X1-old and 7080X1-NMN-before-tumor) (n=5 each). (**N**) Kaplan-Meier survival analysis of 7080X1-bearing mice following intracranial injection of 5×10^5^ cells (*left*, n=7 for 7080X1-old; n=8 for 7080X1-NMN-before-tumor), 5×10^4^ cells (*middle*; n=5 each), and 5×10^3^ cells (*right*; n=5 each).

Given that recent literature has pointed out that boosting the NAD^+^ pathway fuels the growth of glioblastoma cells (*24, 25*), the striking differences between NMN-before-tumor and NMN-after-tumor suggest that the anti-tumor effect by NMN can be achieved by systemic administration only prior to tumor challenge (benefit for prevention) but not after tumor formation (no effect on therapy). To support this interpretation, the NAD^+^ activity measured in these tumors showed a trend of partial reversal (although not reaching statistical significance) of the pathologically activated intra-tumoral NAD^+^ activity in old mice by the systemic NMN treatment prior to tumor challenge (**fig. S9F**).

Similar to tumors in young and old mice subjected to parabiosis, the NMN-before-tumor models showed a rise in mRNAs encoding proteins implicated in the neuronal/synaptic pathways (**Fig. 4F** and **fig. S10, A and B**). A comparison of mRNAs encoding neuroinflammatory genes (i.e. *Ifng*, *Tnfa*, *Nfkb*, *Il6* mRNAs) in ms7080 tumors indicated that NMN pre-treatment dramatically reversed these age-associated elevations; in fact, the abundance of these mRNAs fell below the baseline levels seen in young mice (**fig. S11A**). As expected, expression of mRNAs in the NAD^+^ pathway (*Nampt*, *Cd38*, *Sirt1*, *Parp1*, *Nmnat1-3*, and *Naprt* mRNAs) were largely reversed by NMN pre-treatment for these tumors (**fig. S11B**). The abundance of *Slitrk6* mRNA, as measured by RNA-seq and RT-qPCR analyses, was significantly higher in NMN-before-tumor tissues (**Fig. 4, G and H**). IHC also showed the recovery of the SLITRK6 intensity levels in NMN-before-tumor from markedly diminished levels in the old untreated control (**Fig. 4I**).

Given that NMN pre-treatment conferred survival benefit in these glioblastoma models, we next examined whether the tumors developing in brains from mice pre-treated with NMN had fewer tumor-initiating cells (TICs). To this end, we resected tumors from old mice with and without NMN pre-treatment and established secondary glioma sphere cultures termed 7080X1-old and - NMN-before-tumor, respectively (**Fig. 4J**). The 7080X1-NMN-before-tumor cells exhibited reduced growth in culture and reduced ability to for clonal spheres, as compared to both 7080X1-young and 7080X1-old cells (**fig. S12, A and B**). Similar to the ms7080 tumor counterparts, RT-qPCR analysis indicated that the expression level of *Cd133* mRNA, a surrogate marker of tumor initiation, increased in 7080X1-old cells to levels comparable to those in 7080X1-NMN-before-tumor cells (**fig. S12C**). GO analysis of the RNA-seq data showed that 7080X1-NMN-before-tumor cells were predominantly associated with neuron/synapse-associated pathways (e.g. synapse, generation of neuron, synapse part) (**fig. S12D**). As expected, RT-qPCR analysis from these three cell cultures showed elevated *Slitrk6* mRNA in 7080X1-NMN-before-Tumor cells as compared to 7080X1-old cells (**Fig. 4K**).

We then performed intracranial passaging of these 7080X1 cell lines using young WT mice (**Fig. 4J**). Histopathological examination of these secondary tumors revealed a striking reduction of tumor grade in one-third of the tumors from glioblastoma to Grade III glioma in the 7080X1-NMN-before-tumor line, unlike the 7080X1-old, in which all resultant tumors were glioblastoma with prominent central necrosis and microvascular proliferation (**Fig. 4, L and M, and fig. S12, E and F**). This finding was associated with a significant improvement in survival of mice bearing 7080X1-NMN-before-tumor cells as compared to 7080X1-old tumor cells (**Fig. 4N**). Additionally, *in vivo* serial dilution experiments – the gold standard for measuring TIC subpopulations in each tumor – showed that the mice carrying 7080X1-NMN-before-tumors experienced significantly fewer tumor-burden death as compared to those with 7080X1-old tumors (**Fig. 4N**). Finally, we compared the relative abundance of NAD^+^ and NADH in these 7080X1 models *via* two different culture conditions, termed ‘low NAM’ or normal. We observed significantly higher NAD^+^ and NADH levels in 7080X1-old cells relative to 7080X1-NMN-before-tumor cells, indicating that, similar to the young tumor cells, intratumoral NAD^+^ is lower in 7080X1-NMN-before-tumor cells (**fig. S12G**). Collectively, these data suggest that the survival benefits of NMN pre-treatment for tumor-bearing mice was due at least in part to the diminished TIC subpopulations in tumors.

### Recovered neuronal gene signature in the aging brain by NMN

In light of the phenotypic changes elicited by parabiosis and NMN, we next investigated the cellular effectors activated by the NAD^+^ pathway in old brains (**Fig. 5A**). Following the systemic administration of NMN, we performed RNA-seq analysis of non-tumor-bearing brain samples derived from young, old, Old-para, and NMN-treated old mice (Old-NMN). Heat map clustering analysis showed a change in overall gene expression profiles between these groups (**Fig. 5B**). PCA with PC1 as the predominant determinant as high as 99.7% demonstrated a trend of transcriptomic signature changes from young to old mice which was at least partially reversed both by parabiosis and by NMN pre-treatment, albeit to different degrees among the mice studied (**Fig. 5C**). Similar to the results with the parabiosis models, the aging-induced shift in gene expression signature in cortical tissues was markedly larger than that in striatal tissues (**fig. S12A**). We then separated the genes in the RNA-seq data into those related to the four major cell types in the brain; among them, neuronal genes (*26*) exhibited the most striking shift in the gene signature in cortices of young, old, and old-NMN brains. In contrast, the gene signatures of the other three gene sets (astrocytic, oligodendrocytic, and microglial) did not display a clear trend with aging or NMN treatment (**Fig. 5D**). Ingenuity Pathway Analysis (IPA) of the RNA-seq data for pathways and biological functions confirmed an aging-related reduction of the genes implicated in neurotransmitter metabolism, release of acetylcholine, and behavior pathways, all of which were reversed by NMN pre-treatment of old mice (**Fig. 5E**).

**Figure 5.**
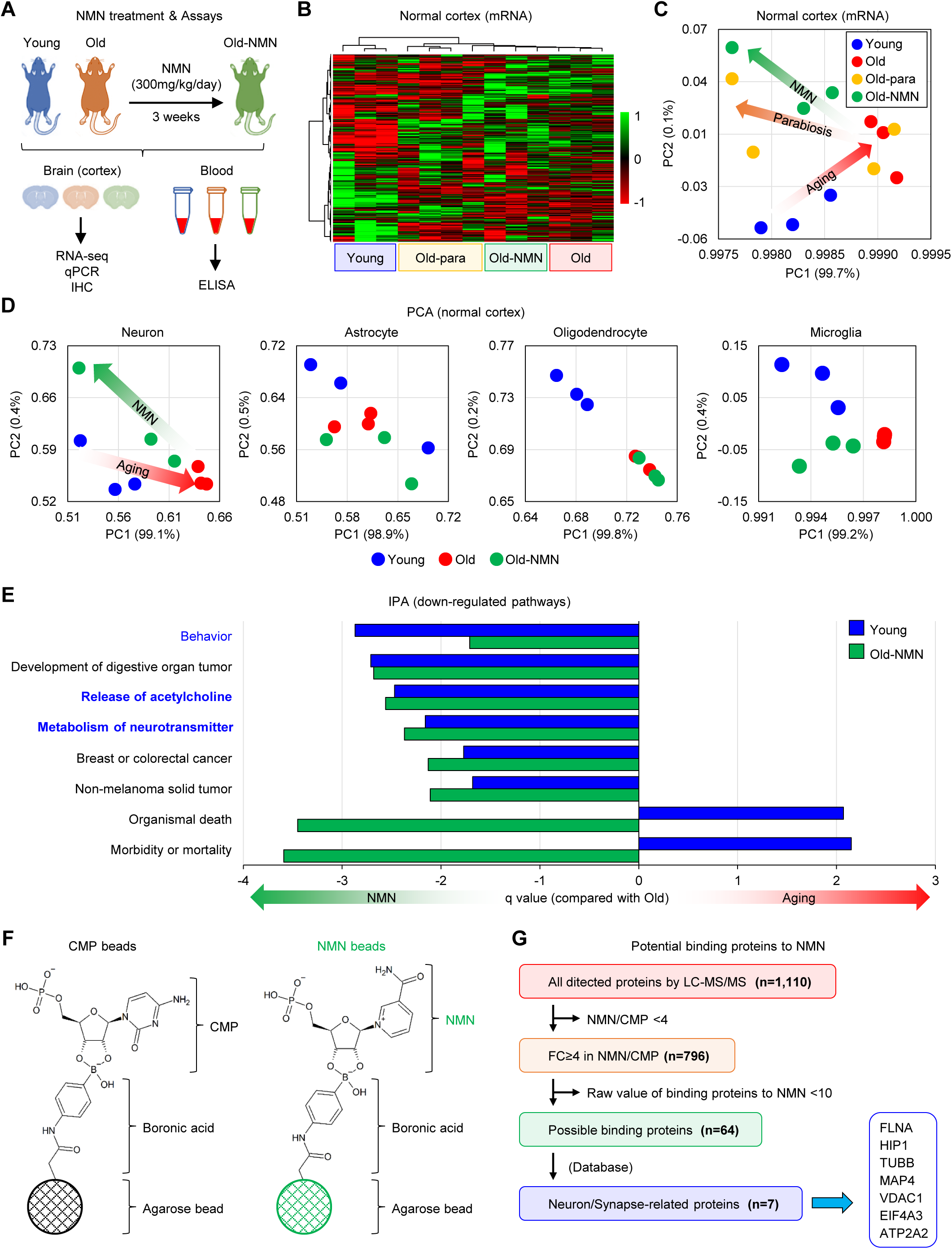
Rejuvenation of neuronal gene signature in normal cortices by NMN treatment. (**A**) Schematic illustration of the timing of NMN administration and sample collections. (**B**) Heat map of RNA-seq data of indicated normal cortical samples (n=3 for young, old, and old-NMN; n=4 for old-para). (**C**) PCA comparing young, old, old-para, and old-NMN mouse cortices. (**D**) PCA comparing genes associated with neurons, astrocytes, oligodendrocytes, and microglia in normal cortices (n=3 for each group). (**E**) Ingenuity pathway analysis of RNA-seq data indicating the activation of specific processes in indicated mouse cortices. (**F**) Schematic of the immobilization of Cytidine 5′-monophosphate (CMP) and NMN on agarose beads. (**G**) Flow chart of the methodology to enrich proteins through the LC-MS/MS experiments with patient-derived glioma spheres.

Additionally, we developed a method to immobilize NMN on beads (**fig. S12B and S12C**). In order to identify proteins binding to NMN, we incubated these beads with membrane proteins and performed LC-MS/MS analysis (**Fig. 5F**). Seven out of 64 proteins identified as associating with NMN were synapse-associated proteins, including FLNA, HIP1, TUBB, MAP4, VDAC1, EIF4A3, and ATP2A2 (**Fig. 5G**). Consistent with the results of the parabiosis models, we confirmed that NMN administration partially recovered NAD^+^ and NADP levels in the blood samples from old mice (**fig. S12D**). These data suggest that NMN might activate cortical neurons through association with neuronal/synaptic membrane proteins to attenuate an aging-related progression toward a tumor-permissive brain microenvironment.

### BDNF contributes to NMN-mediated reprogramming in brains and tumors

To investigate the molecular mechanisms in aging brain to elicit phenotypic changes by NMN, we first examined the expression changes of SLITRK6 in aging normal brains. IHC analysis revealed a striking recovery of SLITRK6 signals in the normal cortex of old NMN-treated mice relative to old untreated cortex, which were comparable to those of young untreated control (**Fig. 6A**). IPA analysis of the RNA-seq data to identify key effectors of the NAD^+^ pathway revealed that brain-derived neurotrophic factor (BDNF) was the most significantly downregulated protein in the old group, and that this trend was largely reversed by NMN treatment (**Fig. 6B**). As expected, a number of inflammatory molecules showed the opposite trend (**Fig. 6B**). A comparative analysis of BDNF interactions revealed downregulation of receptors for BDNF, including NGFR and NTRK1, both in old *vs*. young mice and in old *vs*. old-NMN mice (**Fig. 6C**). Among the four well-studied BDNF exons (exons 1, 2, 4, and 6), NMN induced primarily the recovery of exon 4, previously identified to be excluded with aging and be associated with the NAD^+^ pathway (*27–29*) (**Fig. 6D** and **fig. S13A**).

**Figure 6.**
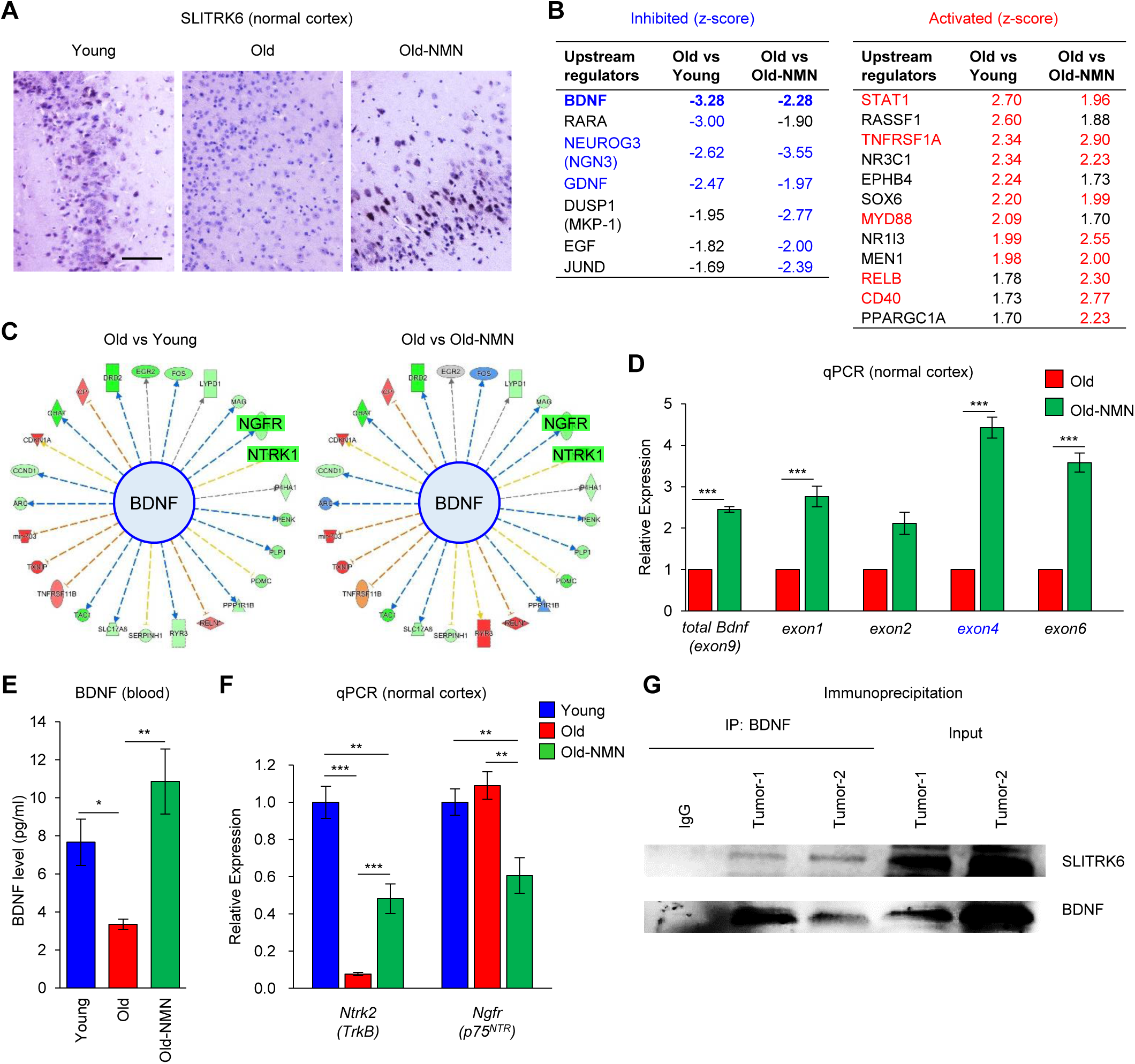
BDNF is a key effector of the response to glioma by the young brain. (**A**) Representative IHC analysis of SLITRK6 levels in normal cortices of young, old, and old-NMN mice. Scale bar represents 100 µm. (**B**) List of inhibitory and activating regulators identified by RNA-seq analysis with normal cortices in young, old, and old-NMN mice. (**C**) Function map of BDNF downstream genes regulated in old cotices compared to young cortices (left) and to old-NMN cortices (right). Red indicates upregulated and green indicates downregulated mRNAs. The shape corresponds to the significance. (**D**) RT-qPCR analysis of the levels of total *BDNF* mRNA as well as *BDNF* exons 1, 2, 4, and 6 in old and old-NMN cortices. Data are the means ± SD (n=6). \*\*\**p*<0.001. (**E**) ELISA analysis of BDNF levels in mouse blood; data are the means ± SD (n=4). \**p*<0.05, \*\**p*<0.01. (**F**) RT-qPCR analysis of the levels of *Ntrk2* (*TrkB*) and *Ngfr* (*p75^NTR^*) mRNAs in young, old, and old-NMN cortices. Data are the means ± SD (n=6 each). \*\**p*<0.01, \*\*\**p*<0.001. (**E**) RT-qPCR analysis of *Nfkb1*, *Nfkb2*, and *Ccl2* mRNAs, encoding inflammatory markers, in young, old, and old-NMN cortex. Data are the means ± SD (n=6). \*\*\**p*<0.001. (**G**) Immunoprecipitation (IP) analysis with anti-BDNF antibody followed by immunoblot for SLITRK6 in ms7080 tumors. Input tumor lysates and IgG IP were included as positive and negative controls, respectively.

In addition to these changes in BDNF levels in the cerebral cortex, the aging-associated reduction of the systemic levels of circulating BDNF in the blood was totally rescued by NMN treatment (**Fig. 6E**). Furthermore, NMN administration reversed the aging-related changes in *Ntrk2* (transcript for TrkB), encoding a receptor of mature BDNF (mBDNF) and a known agonist for neurons, and in *Ngfr* (transcript for p75^NTR^), encoding a receptor of precursor of BDNF (proBDNF) and a known antagonist for neurons (**Fig. 6F**). As anticipated, the expression levels of neuroinflammatory markers (*Nfkb1*, *Nfkb2*, and *Ccl2* mRNAs) in these samples displayed opposite patterns, as determined by RT-qPCR validation of IPA data (**fig. S13B**). The ensuing analysis of a possible physical interaction of BDNF and SLITFK6 proteins by immunoprecipitation followed by Western blotting analysis demonstrated an association between the two proteins (**Fig. 6G**). Collectively, these data suggest a novel molecular signaling axis whereby BDNF forms a protein complex with SLITRK6 linked to the development of a cortical aging, tumor-favorable microenvironment.

### Reduction in NMN-regulated BDNF aggravates tumor phenotype in old brains

Next, we sought to determine whether the aging-associated glioblastoma malignancy is regulated by NAD^+^ and BDNF by performing tumor challenges in BDNF^+/-^ mice with and without NMN pre-treatment (**Fig. 7A**). The tumors in BDNF^+/-^ mice were histopathologically more malignant, harboring highly condensed tumor cells with larger central areas of necrosis, and the mice had shorter survival than tumor-bearing WT mice (**Fig. 7B and C**). Importantly, the survival benefit of mice in the NMN pre-treatment group was absent in BDNF^+/-^ underscoring the role of BDNF in the anti-tumor effect of NMN (**Fig. 7C**). These data suggested that the reversal of the tumor permissive microenvironment activated by NAD^+^ in older mice is primarily regulated by the circulating BDNF.

**Figure 7.**
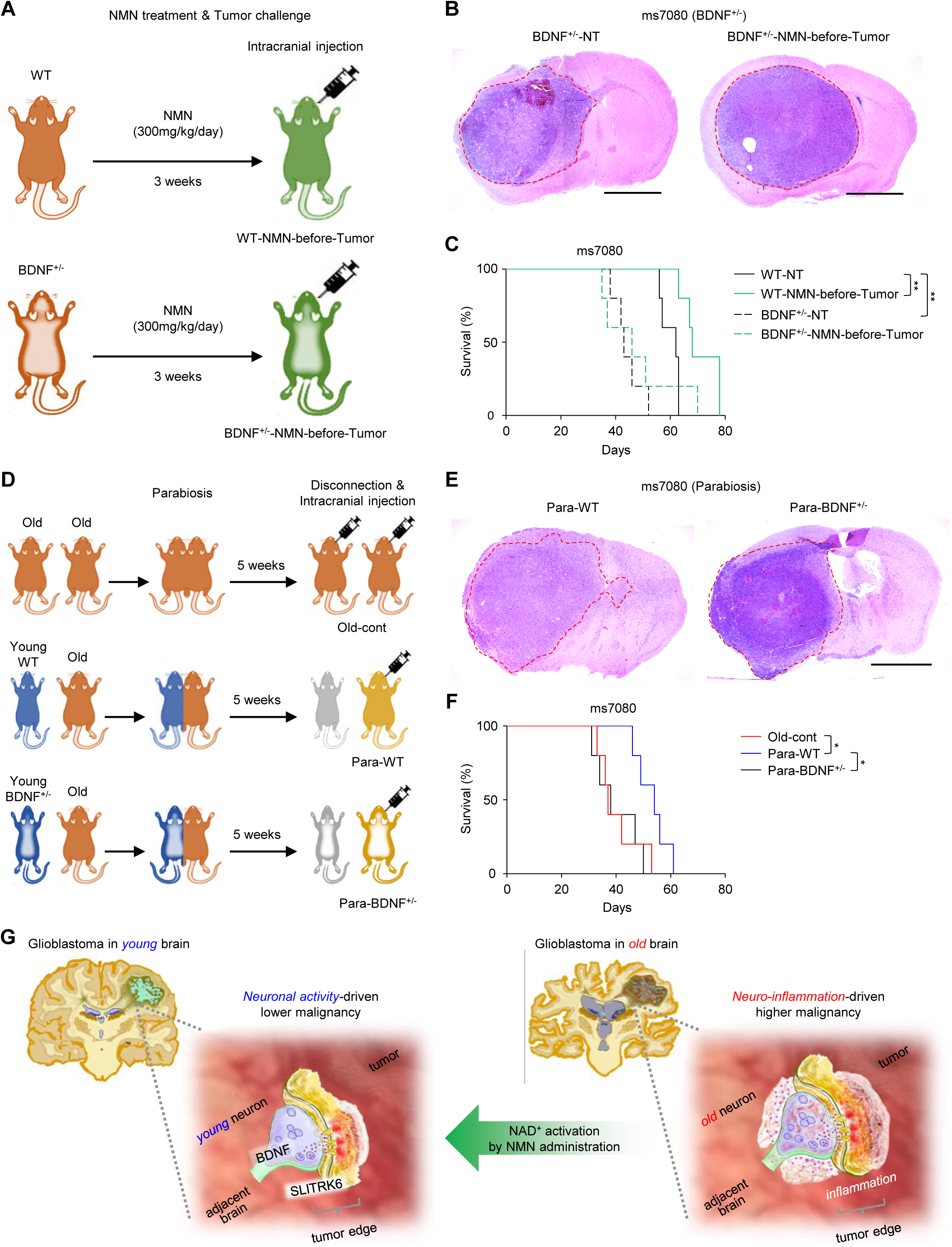
Decline in BDNF levels contributes to aggravating tumor phenotype in old brains. (**A**) Schematic of the timing of NMN administration together with ms7080 challenge in wild-type (WT) and BDNF^+/-^ mice. (**B**) H&E staining analysis of mouse brains bearing m7080 tumors in BDNF^+/-^ mice with and without NMN pre-treatment. BDNF^+/-^-NT indicates without NMN pre-treatment. Scale bars represent 2 mm. (**C**) Kaplan-Meier survival analysis of WT (n=5 for NT and NMN-before-tumor) and BDNF^+/-^ (n=5 for NT and NMN-before-tumor) mice bearing ms7080 tumors following NMN pre-treatment, compared to control mice. (**D**) Schematic of the chronological order for parabiosis of old mice with old, young WT, or BDNF^+/-^ mice together with ms7080 injection. (**E**) H&E staining analysis of brains in Para-WT mice (left), compared to Para-BDNF^+/-^ mice (right). Scale bar represents 2 mm. (**F**) Kaplan-Meier survival analysis of the indicated mice with parabiosis (n=5 for each). (**G**) Schematic of the NAD^+^-BDNF-mediated signaling hypothesis in young and old glioblastomas.

Lastly, we asked whether systemically circulating BDNF, and not only the brain-resident BDNF, determines the aging-associated anti-tumor effect of BDNF in the brain. To this end, we performed parabiosis followed by tumor challenge with Old-cont, Old-para (termed ‘Para-WT’), and old mice receiving young BDNF^+/-^ mouse blood (termed ‘Para-BDNF^+/-’^) (**Fig. 7D**). Strikingly, the m7080-derived tumor phenotype of Para-BDNF^+/-^ mice displayed a clear difference from Para-WT, exhibiting densely packed tumor lesions with markedly larger central necrosis – similar to the lesions seen in Old-cont (**Fig. 7E**). As a result, the young blood-engendered survival benefit for older mice almost completely disappeared when the blood was received from young BDNF^+/-^ mice (**Fig. 7F**). Collectively, these findings indicate that the decline of circulating BDNF is a major effector of the tumor-favorable environment seen in old mouse brain, triggered by the age-associated reduction in NAD^+^ (**Fig. 7G**).

## Discussion

Aging accelerates the risk of cancer and aggravates malignant phenotypes. This study uncovered that the aging-induced cancer-permissive changes in the brain microenvironment can be reversed, at least in part, by systemic factors, as confirmed by both parabiosis between young and old animals and systemic administration of the NAD^+^ pathway booster NMN. This observation, if applicable to humans, underscores the potential of using NMN (and other means of activating the NAD^+^ pathway) to reduce the risk of cancer formation and deterioration through rejuvenation of the brain and other organs. It remains unclear why blending blood circulation between young and old mice for as long as 5 weeks (approximately 7-9 years for humans) almost completely flipped the tumor phenotypes between them, rather than rendering intermediate phenotypes (e.g. tumors corresponding to “middle-aged brains”). These data may reflect the existence of unknown factors that trigger opposite phenotypes (young converted to old-like and old converted to young-like). Further studies are required to answer this question.

Older glioblastoma patients often survive less than a year from the time of diagnosis. This aging-driven elevation of malignancy is thought to be closely linked to neuroinflammation, although the possibility remains that (hyper)active neurons may aggravate glioblastomas pathoetiology even in older patients. The mechanism whereby neuroinflammation is initiated and promoted in glioblastoma needs further investigation. Inflammation is generally stimulated by soluble factors (e.g. chemokines, cytokines, metabolites) secreted by either resident cells such as microglia and astrocytes or by cells external to brain tissues, such as macrophages. Identifying the specific cellular factors responsible for aging-mediated neuroinflammation and tumor-permissive microenvironment would be very helpful in designing therapies to combat the effect of aging on cancer phenotype. Cortical neurons are terminally differentiated non-cycling cells, yet recent studies have shown that they develop a senescent-like status through accumulation of age-related DNA damage. Senescent cells are known to secrete powerful paracrine factors, including proinflammatory cytokines that spread to neighboring cells. Whether this senescent program caused by aged neurons helps to orchestrate the deteriorating microenvironment in aged tumors warrants future study.

Both parabiosis with younger mice and systemic NMN administration resulted in notably similar tumor-preventive changes in the old-brain microenvironment. In the brain, NMN protects neurons from chemotherapy-induced degeneration and cell death after stroke(*30, 31*). Our recent study revealed that NMN pre-treatment of CD38 (NADase) KO mice followed by ms7080 tumor challenge improved glioblastoma malignancy and altered intra-tumoral NAD^+^ activity, supporting the notion that the action of systemic NMN administration directly controls NAD^+^ activity in murine glioblastoma. Nonetheless, this earlier work did not investigate the possible change in the BDNF signaling in the brain or tumors. In the present study, as illustrated by the remarkable difference between prevention and therapy for NMN in mice with tumor challenge, NMN treatment appears to be a double-edged sword, providing nutrients for normal somatic cells that is also highjacked by cancer cells as an energy source (*24*). Accordingly, plans to boost the NAD^+^ pathway only in non-cancerous cells must include the shutting off of the cancer cell-specific receptor for NMN. Recently, SLC12a8 was identified as a transporter of NMN in the intestine, but this transporter is not expressed in the brain or glioblastoma cells(*32*). This study identified a set of (brain) cancer-specific transporter candidates for NMN, although validation awaits. Through future investigation, one may enable the use of NMN for not only prevention but also treatment of glioblastoma, particularly in old patients.

Based on the two sets of data using the BDNF^+/-^ mice combined with pre-treatment of NMN or parabiosis, BDNF is likely a *bona fide* gate-keeper for NMN-driven NAD^+^ activity in the brain. However, caution is warranted, as BDNF may display antagonistic pleiotropy (possibly by forming a protein complex with SLITRK6) to provoke a fundamental tumor-permissive microenvironmental architecture, particularly in a young adult-specific manner. How the NAD^+^ pathway regulates BDNF secretion and activity in the brain microenvironment needs additional study. Identification of SLITRK6 as a novel binding protein for BDNF raises a possibility that this protein complex may communicate signals derived from the elevated NAD^+^ activity to rejuvenate the tumor phenotype.

In conclusion, this study has developed a new NAB subclassification based on novel gene expression patterns related to neuronal activity and identifying clinically relevant subtypes. Furthermore, we performed a set of *in vivo* experiments with parabiosis and NMN administration using 4 mouse glioblastoma tumor models and 1 human patient-derived tumor model for tumor challenge in the brain. These experiments uncovered effects of aging and rejuvenation on tumorigenesis. The gene signature in both paradigms exhibited an aging-associated shift from neuronal activity to neuroinflammation, accompanied by substantial elevation in tumor aggressiveness. This aging-associated molecular and phenotypic changes can be reversed by either receiving young blood or NMN administration, underscoring the plasticity of the brain in susceptibility to disease. Validation of these interventions in human glioblastoma can pave the way towards novel age-dependent cancer therapy regimens.

## Acknowledgements

We would like to express our sincere appreciation to all the patients and families, who kindly allowed us to obtain their tumor samples for this study. We would also thank all our collaborating scientists, as well as the assigned reviewers and Editor for this manuscript, for the constructive comments and suggestions. We acknowledge the contribution by all the members in the Nakano and Gorospe laboratories (past and present) for technical help. We also thank Oriental Yeast Co., Ltd. for supply of NMN, G. Coppola for intellectual input on this project, L. Chow for supply of AR006 neurosphere, and the rest of our team members for their valuable suggestions. This work was supported by NIH grants R01NS083767, R01NS087913, R01CA183991, and R01CA201402 to IN and by the NIA IRP (NIH) to MG and RM.

## Author Contributions

Leading conceptualization of the study: IN. Financial support: IN, MG. Overall design of the study: DY, IN, with input from TK, DH, HIK, TS, MG. Laboratory practice: DY, VLF, KS, SO, SS, SB, MSP, SY, JRH. Analysis of data; DY, RBM, KS, RK, MG, IN. Establishment of NSCL61 cells: TK. Drafting the article: DY, MAN, IN. Critical revision of the article: MG, IN. All authors had substantial input to the logistics of the work and revised and approved the final manuscript. The authors know their accountability for all aspects of the study ensuring that questions regarding the accuracy and integrity of any part are appropriately investigated and resolved. The corresponding author had full access to all of the data and the final responsibility to submit the publication.

## Materials and Methods

### Patients, Specimens, and Ethics

For the pre-clinical studies, the previously characterized patient-derived glioma sphere model (1051X1) established and described elsewhere (*33*) was used. In short, with signed patient consent, the senior author (IN) performed supra-total resection of glioblastoma tumors under the awake setting and resected tumor to achieve maximal tumor eradication without causing any permanent major deficit in the patients. After confirmation of enough tumor tissue was secured for the clinical diagnosis, the remaining tissue was provided to the corresponding scientists following de-identification of the patient information. This patient-derived glioma model was periodically checked for mycoplasma using the Short Tandem Repeat (STR) analysis. All pre-clinical work was performed under an Institutional Review Board (IRB)-approved protocol (N150219008) compliant with guidelines set forth by the National Institutes of Health (NIH).

### Cell culture

Murine glioma-derived (neuro)sphere cultures were established from tumors formed by *Cre; Nf1*^f/+^; *p53*^f/f^; *Pten*^f/+^ mice (ms7080)(*19*), *CreER; Pten*^loxP/loxP^; *Tp53*^loxP/loxP^; *Rb1*^loxP/loxP^ mice (AR006)(*20*), *Cre; Nf1*^f/f^; *p53*^f/f^ mice (ms6835)(*21*), and by transforming p53-deficient neural stem cells (NSC) with oncogenic HRas^L61^ (NSCL61)(*22*). Characterization of the patient-derived 1051 spheres is described in our prior study (*33*). Both murine and patient-derived glioma spheres were cultured in defined medium containing DMEM/F12/ Glutamax (Invitrogen) supplemented with B27 (Miltenyi Biotec), heparin (2.5 mg/mL), basic fibroblast growth factor (bFGF) (Peprotech, 20 ng/mL), and epidermal growth factor (EGF) (Peprotech, 20 ng/mL). Growth factors (bFGF and EGF) were added twice a week and the culture medium was changed every 7 days.

### RNA isolation and Quantitative Real-Time PCR (RT-qPCR) analysis

Total RNA was extracted using the RNeasy mini kit (QIAGEN) according to the manufacturer’s instructions. RNA concentration was determined using Nanodrop One (Thermo Scientific). cDNAs were synthesized using iScript reverse transcription supermix (Bio-Rad) according to the manufacturer’s protocol. RT-qPCR analysis was performed on StepOnePlus thermal cycler (Thermo scientific) with SYBR Select Master Mix (Thermo scientific). *GAPDH*/*Gapdh* mRNA was used as an internal control. Primer sequences are shown in Supplementary Table 1.

### Western blotting

Cells were lysed for 30 min on ice in RIPA buffer (Sigma) containing 1% protease and 1% phosphatase inhibitor cocktails (Sigma). Protein samples were quantified using the Bradford assay reagent (Bio-Rad) according to the manufacturer’s instructions. The proteins were transferred onto a PVDF membrane and the membranes were probed overnight at 4 °C with the appropriate primary antibodies recognizing SLITRK6 (Invitrogen) or β-Actin (ACTB, Cell Signaling). After incubation with HRP-conjugated second antibodies, staining was visualized with Amersham ECL Western Blot System (GE Healthcare) and images were obtained with ImageQuant LAS 500 (GE Healthcare).

### Immunoprecipitation (IP)

Brain tissues were collected and lysed in IP lysis buffer (Pierce) supplemented with protease inhibitors, incubated on ice for 15 min, and cleared by centrifugation at 13,000 rpm at 4°C for 15 min. After a preclearing step with protein A/G-agarose beads (Upstate), protein lysate (1 mg) was subjected to IP (overnight at 4°C) in the presence of beads carrying antibodies that recognized BDNF (LSBio), SLITRK6 (Invitrogen), or with isotype control antibodies.

### Immunohistochemistry (IHC)

Human glioma tissues (n=20) were collected at the Ehime University after obtaining the written informed consent forms from the patients. IHC was performed as previously described and signals were detected using the DAB substrate kit (Vector) (*33–35*). For double staining, donkey IgG H&L (alkaline phosphatase) pre-adsorbed antibody (Abcam) was used to detect primary antibodies and detected by the liquid fast-red substrate kit (abcam). Primary antibodies used in this study recognized SLITRK6 (Invitrogen).

### Animal experiments

All animal experiments were performed at the UAB under an Institutional Animal Care and Use Committee (IACUC)-approved protocol according to NIH guidelines. C57BL/6 mice and immunocompromised mice (SCID Beige and NSG) were purchased from Charles River and Jackson Laboratory. For intracranial injection, mice were anesthetized with ketamine/xylazine and fixed in place and dissociated BC cells were stereotactically injected into the striatum of mice as described (*33–35*). Mice were placed on a stage warmed at 37°C until they were fully awake.

### Parabiosis

Female mice were placed in a cage for two weeks to assure peaceful cohabitation. Two animals were anesthetized using 1-5 to 2% isoflurane, shaved at approximately 1 cm above the elbow to 1 cm below the knee, prepared by thoroughly wiping (3x) with Betadine-soaked wipes followed by alcohol wipes, and placed on a heated pad covered by a sterile pad. For analgesia, Carprofen and Buprenorphine were administered subcutaneously at a dose of 10 mg/kg and 0.1 mg/kg, respectively. Longitudinal skin incisions were performed to the shaved sides of each animal starting at 0.5 cm above the elbow all the way to 0.5 cm below the knee joint. Following the incision, the skin was gently detached from the subcutaneous fascia by holding if up with a pair of curved forceps. The fascia was then separated with a second pair to create 0.5 cm of free skin. The left olecranon of one animal will be joined to the right olecranon of the other. Both olecranons and knee joints were clearly distinguishable following the skin incision. To facilitate the joining, the elbow of the first mouse was bent and a needle of the non-absorbable 3-0 suture will be passed under the olecranon. Following the attachment of the joints, the skin of the two animals was joined with continuous absorbable 5-0 Vicryl suture starting ventrally from the elbow towards the knee. To prevent skin rupture and separation tight suture closure of the skin in the area around the elbows and knees was performed. To prevent dehydration, 0.5 mL of 0.9% NaCl was administered subcutaneously to each mouse, and animals were kept on a heated pad until recovery. Following recovery, analgesics carprofen and buprenorphine were given bysubcutaneous injection every 12 hr for 48 hr at the same doses described above. Animals were monitored for signs of pain and given Sulfamethoxazole/Trimethoprim in drinking water (2 mg sulfa/mL +0.4 mg trim/mL) for 10 days.

### NMN Administration

Water consumption was measured prior to the start of NMN administration. NMN was administered in drinking water at 300 mg/kg/day, based on the previously measured water consumption. The NMN solution was prepared twice weekly by dissolving NMN into autoclaved water at the respective doses and sterilized by filtering. Water bottles and cages were changed twice weekly.

### Immobilization of NMN and CMP on agarose beads

Two hundred μL of m-Aminophenylboronic acid–Agarose beads (Sigma) were washed 3 times with 1 mL of binding buffer (200 mM NH4Ac, 30 mM MgCl2, pH 8,9) and incubated with 1 mL of NMN (Oriental Yeast Co., Ltd.) or CMP (Sigma) solution (5 g/l in binding buffer) for 1 hour at room temperature. Remaining concentration of NMN and CMP in the solution were monitored using NanoDropONE spectrophotometer (Thermo Fisher) by absorbance at λ = 270 nm. Beads were washed 3 times with binding buffer and incubated with 1 mL of 2:1 mixture of binding buffer and solubilized membrane proteins purified from glioblastoma cells. After overnight incubation at +4°C with constant agitation, beads were washed once with 2:1 mixture of binding buffer and solubilization buffer (Thermo Fisher) and 4 times with binding buffer. After the last wash NMN, CMP and bound proteins were eluted with 100 μL of 25 mM HCl. Immediately after elution, the eluate was neutralized with 1 M Tris buffer pH 9. Eluted proteins were subjected to LC-MS/MS analysis, performed on a TripleTOF 5600+ mass-spectrometer with a NanoSpray III ion source (ABSciex) coupled with a NanoLC Ultra 2D+ nano-HPLC system (Eksigent) as described previously(*35*). Proteins eluted from empty beads (without NMN or CMP) were used as a negative control.

### ELISA

The concentration of BDNF was measured by using the BDNF ELISA kit (LSbio) according to the manufacturer’s protocol.

### Neurosphere formation assay

Murine GBM neurospheres were seeded into 96-well plates at 1, 10, 20, 30, 40, and 50 cells per well. After 3 days, the numbers of spheres with diameters greater than 60 mm were counted. Data were analyzed as described previously (http://bioinf.wehi.edu.au/software/elda/).

### RNA sequencing

Isolated RNA samples were sequenced commercially at QUICK BIOLOGY (http://www.quickbiology.com). Following depletion of ribosomal (r)RNA, libraries were prepared and sequenced using an Illumina HiSeq 4000 instrument, PE150, for a total of 80 million reads per sample.

### RNA-Seq analysis

STAR (version 2.5.3a) was used to align the raw RNA-Seq fastq reads to the mouse reference genome (GRCm38 p4, Release M11) from Gencode with parameters --outReadsUnmapped Fastx; --outSAMtype BAM SortedByCoordinate; --outSAMAttributes All (*36*). Following alignment, HTSeq-count (version 0.9.1) was used to count the number of reads mapping to each gene with parameters -m union; -r pos; -t exon; -i gene_id; -a 10; -s no; -f bam (*37*). Normalization and differential expression were then applied to the count files using DESeq2.

### Systems Biology analysis

For generating networks, a data set containing gene identifiers and corresponding expression values was uploaded into Ingenuity Pathway Analysis. Each identifier was mapped to its corresponding object in Ingenuity’s Knowledge Base. A fold change cutoff of ±2 and p-value < 0.05 was set to identify molecules whose expression was significantly differentially regulated.

These molecules, called network-eligible molecules, were overlaid onto a global molecular network developed from information contained in Ingenuity’s Knowledge Base. Networks of network eligible molecules were then algorithmically generated based on their connectivity. The functional analysis identified the biological functions and/or diseases that were most significant to the entire data set. Molecules from the dataset that met the fold change cutoff of ±2 and p-value < 0.05 and were associated with biological functions and/or diseases in Ingenuity’s Knowledge Base were considered for the analysis. Right-tailed Fisher’s exact test was used to calculate a p-value determining the probability that each biological function and/or disease assigned to that data set is due to chance alone.

### Gene Expression Data Analysis

Gene Set Enrichment Analysis (GSEA) was performed using available online software (http://software.broadinstitute.org/gsea/index.jsp). Gene Ontology (GO) enrichment analysis also was performed using available website (http://geneontology.org/).

### Measurement of NAD metabolites

Twenty microliters of mouse blood were mixed with 180 μl of 80% methanol and shaken at 1200 rpm for 10 min at 37°C. After centrifugation at 16000 x g for 5 min at 25°C, 100 μl of supernatant were collected and mixed with 40 μl of purified water and 72 μl of chloroform, followed by a vortex mixing for 5 seconds. After centrifugation at 16000 x g for 5 min at 25°C, 50 μl of supernatant were collected and dried in a centrifugal evaporator (CVE-3100, Tokyo Rikakikai Co. Ltd.). For brain analysis, the dissected cerebral cortex was homogenized in ice-cold 80% methanol (tissue:solvent=1:4, w/v) using a beads homogenizer (μT-12, Taitec) at 3200 rpm for 30 seconds two times. After centrifugation at 16000 x g for 30 min at 25°C, 100 μl of supernatant were treated in the same way as the blood samples. Each dried sample was dissolved in 50 μl of 0.1% formic acid in water and 3 μl of the sample solution was injected into to the LC-MS system. Metabolites were separated on a Shim-pack GIST C18-AQ column (3 μm, 150 mm × 2.1 mm id, Shimadzu GLC) with a Nexera UHPLC system (Shimadzu). The mobile phase consisted of 0.1% formic acid in water (A) and 0.1% formic acid in acetonitrile (B). The gradient program was as follows: 0-3 min, 0% B; 3-15 min, linear gradient to 60% B; 15-17.5 min, 95%B; 17.5-20.0 min, linear gradient to 0% B; hold for 4 min; flow rate, 0.2 ml/min. The LC system was coupled with a triple-quadruple mass spectrometer LCMS-8040 or LCMS-8060(Shimadzu). Mass spectrometers were operated with the ESI in positive (for NAM and NMN) and negative ion mode (for NAD and NADP). Ion transitions for multiple reaction monitoring were as follows: NAD, *m/z* 662.10>540.05; NADP, *m/z* 742.00>619.95; NAM, *m/z* 123.10>80.05, NMN, *m/z* 335.05>123.10.

### Statistics

The required sample sizes were estimated on the basis of our previous experience with similar experiments. Statistical analyses and graph generation were performed using XLSTAT 2018.5, SPSS statistical package version 25, and Graphpad Prism 7.0 software. All data were presented as the mean ± SD. P-values <0.05 were considered statistically significant. Statistical differences were determined using unpaired, two-tailed Student’s *t* test. Statistically significant differences in Kaplan-Meier survival curves were determined by log-rank analysis.

### Data and Code availability

Data are available in the Gene Expression Omnibus (GEO) under accession GSE135062 and GSE135210. All custom code used in this work is available from the corresponding author upon request.

## Supplementary figure legends

**Figure S1 (complements Figure 1).**
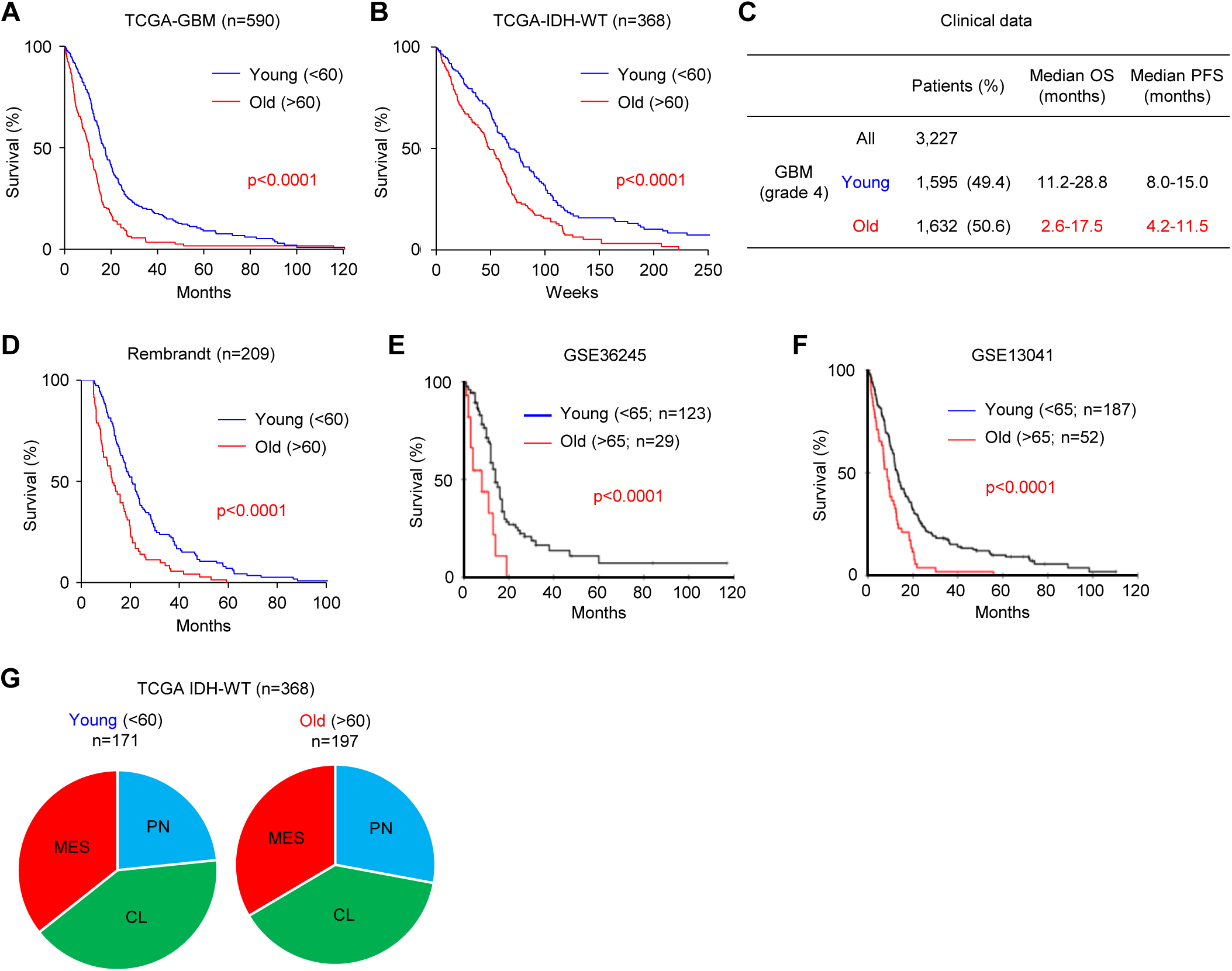
(**A-B**) Kaplan-Meier curve comparing overall survival (OS) of young and old glioblastoma patients in the TCGA-GBM (**A**) and TCGA-WT (**B**) datasets (n= 590 and 368, respectively). *p*<0.0001. (**C**) Table comparing OS and progression-free survival (PFS) in WHO Grade III astrocytomas, as well as young and old glioblastoma patients. (**D-F**) Kaplan-Meier curve comparing OS of young and old glioblastoma patients in the Rembrandt (**D**), GSE36245 (**E**), and GSE13041 (**F**) datasets. (**G**) Pie chart comparing the frequency of glioblastoma molecular subtypes in young and old glioblastoma patients in the TCGA IDH-WT dataset.

**Figure S2 (complements Figure 1).**
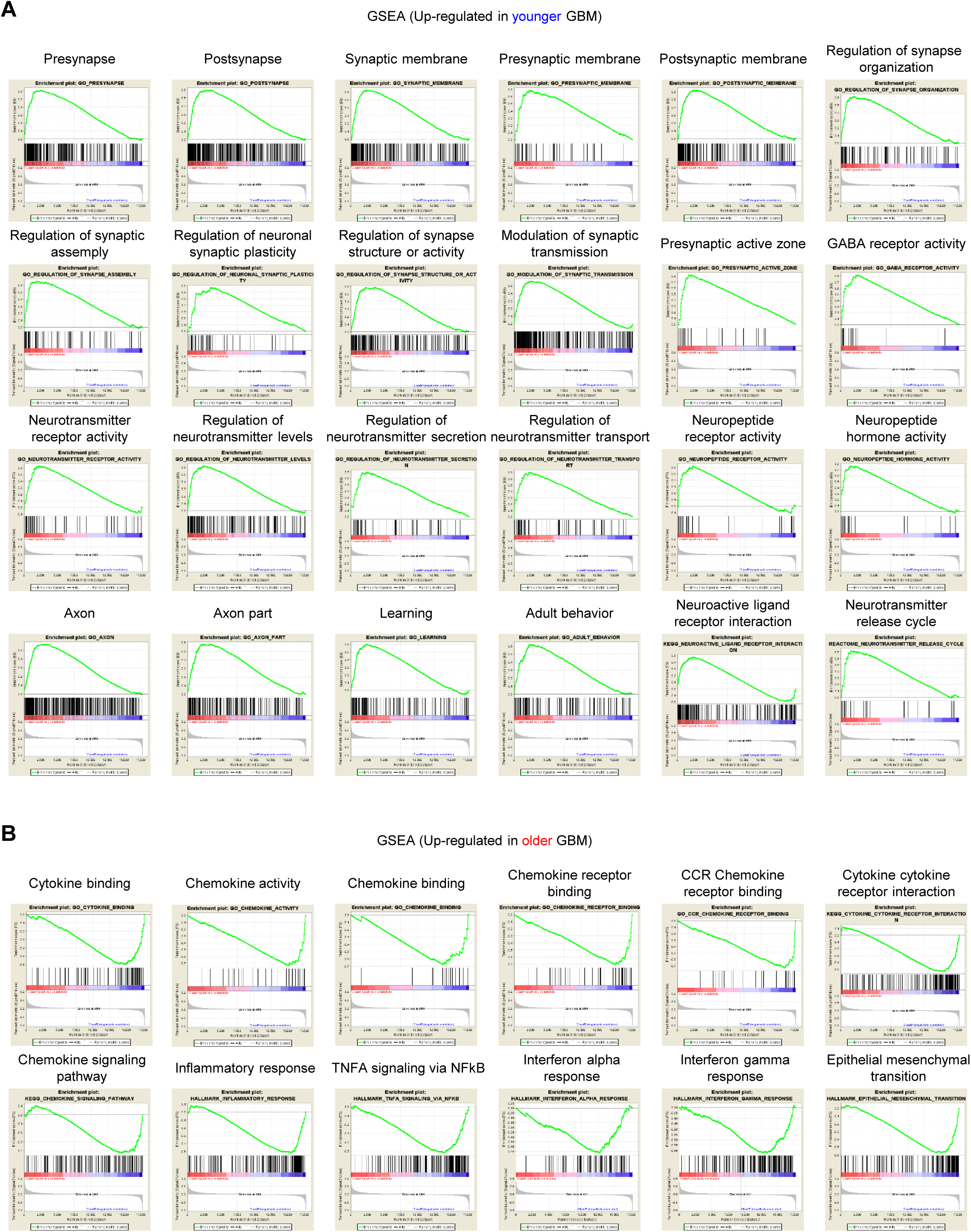
(**A**) GSEA plots of genes associated with neuronal/synaptic pathways in young glioblastoma patients. Gene expression data are obtained from the TCGA database. (**B**) GSEA plots of genes implicated in inflammation in old glioblastoma patients. Gene expression profile data are obtained from the TCGA database.

**Figure S3 (complements Figure 1).**
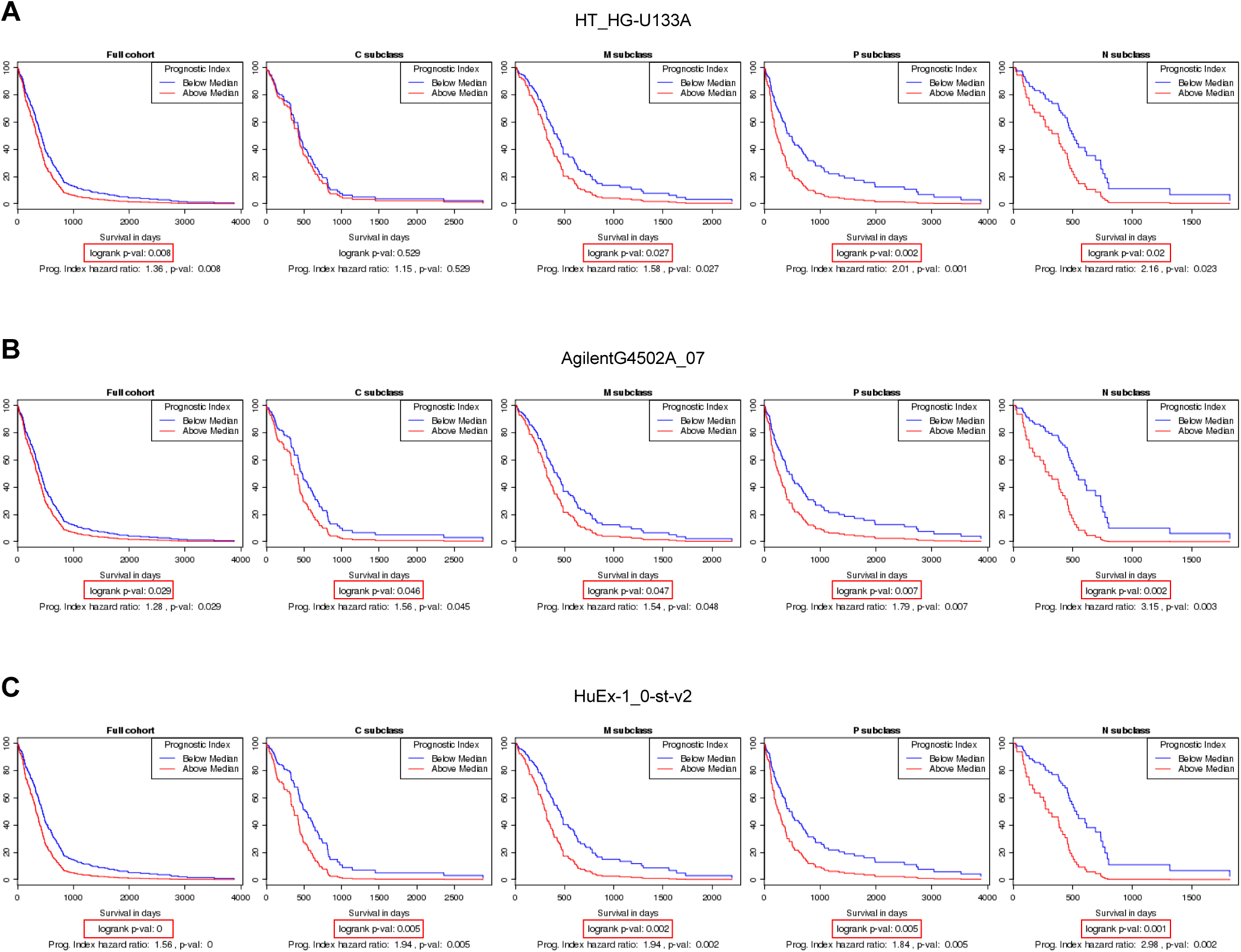
(**A-C**) Kaplan-Meier survival analysis comparing OS of glioblastoma patients with high expression of 11 neuron/synapse associated genes in the HT_HG-U133A (**A**), AgilentG4502A_07 (**B**), and HuEx-1_0-st-v2 (**C**) datasets.

**Figure S4 (complements Figure 1).**
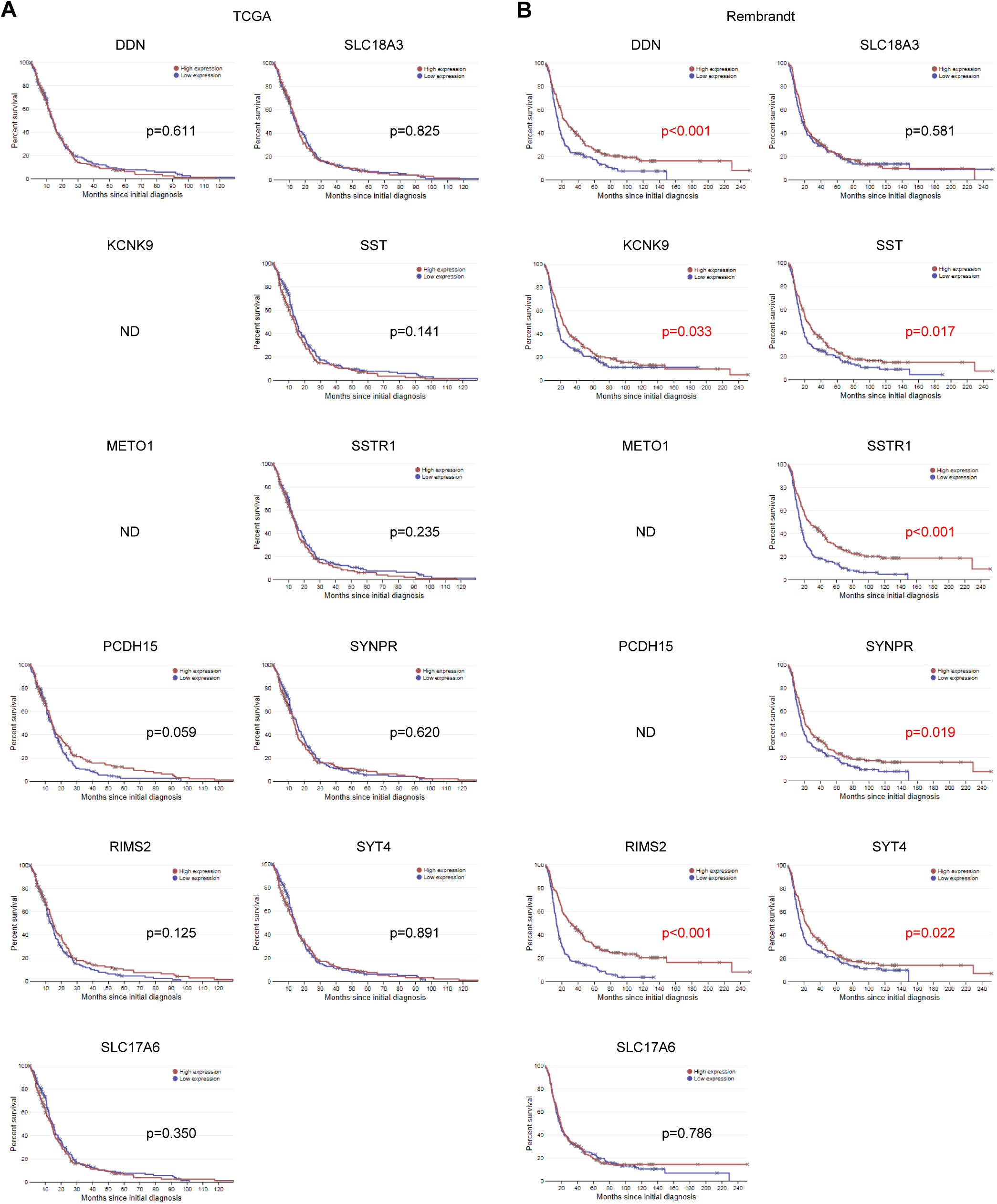
(**A-B**) Kaplan-Meier survival analysis comparing OS of glioblastoma patients based on the levels of 11 neuron/synapse-associated genes in the TCGA (**A**) and Rembrandt (**B**) datasets.

**Figure S5 (complements Figure 2).**
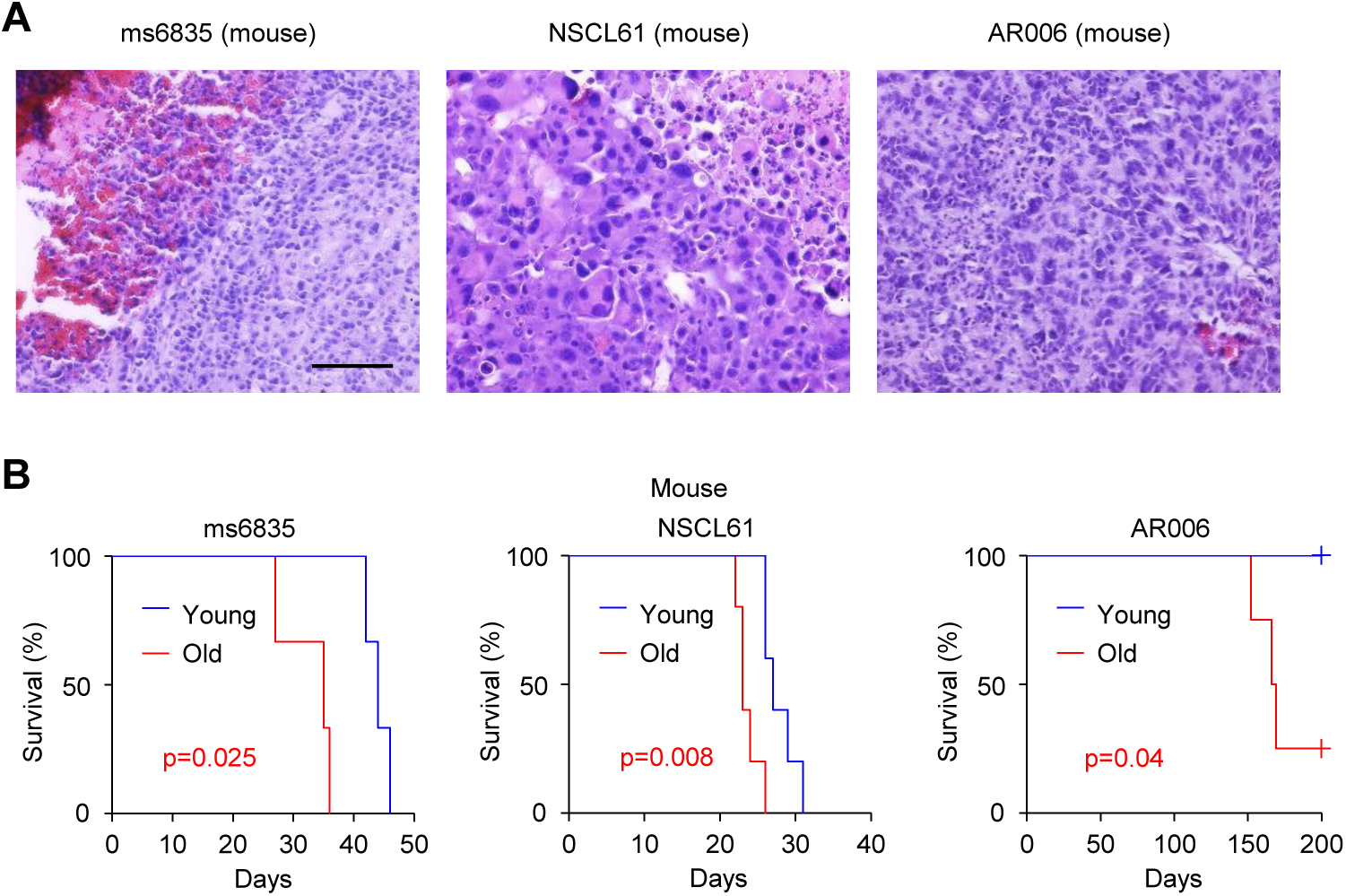
(**A**) Representative H&E staining analysis of mouse brains following intracranial injection of ms6835 (left), NSCL61 (middle), or AR006 (right) cells. Scale bar represents 100 µm. (**B**) Kaplan-Meier curve comparing young and old mice following intracranial injection of mouse ms6835 (left; n=3 for each), NSCL61 (middle; n=5 for each), or AR006 cells (right; n=4 for each).

**Figure S6 (complements Figure 2).**
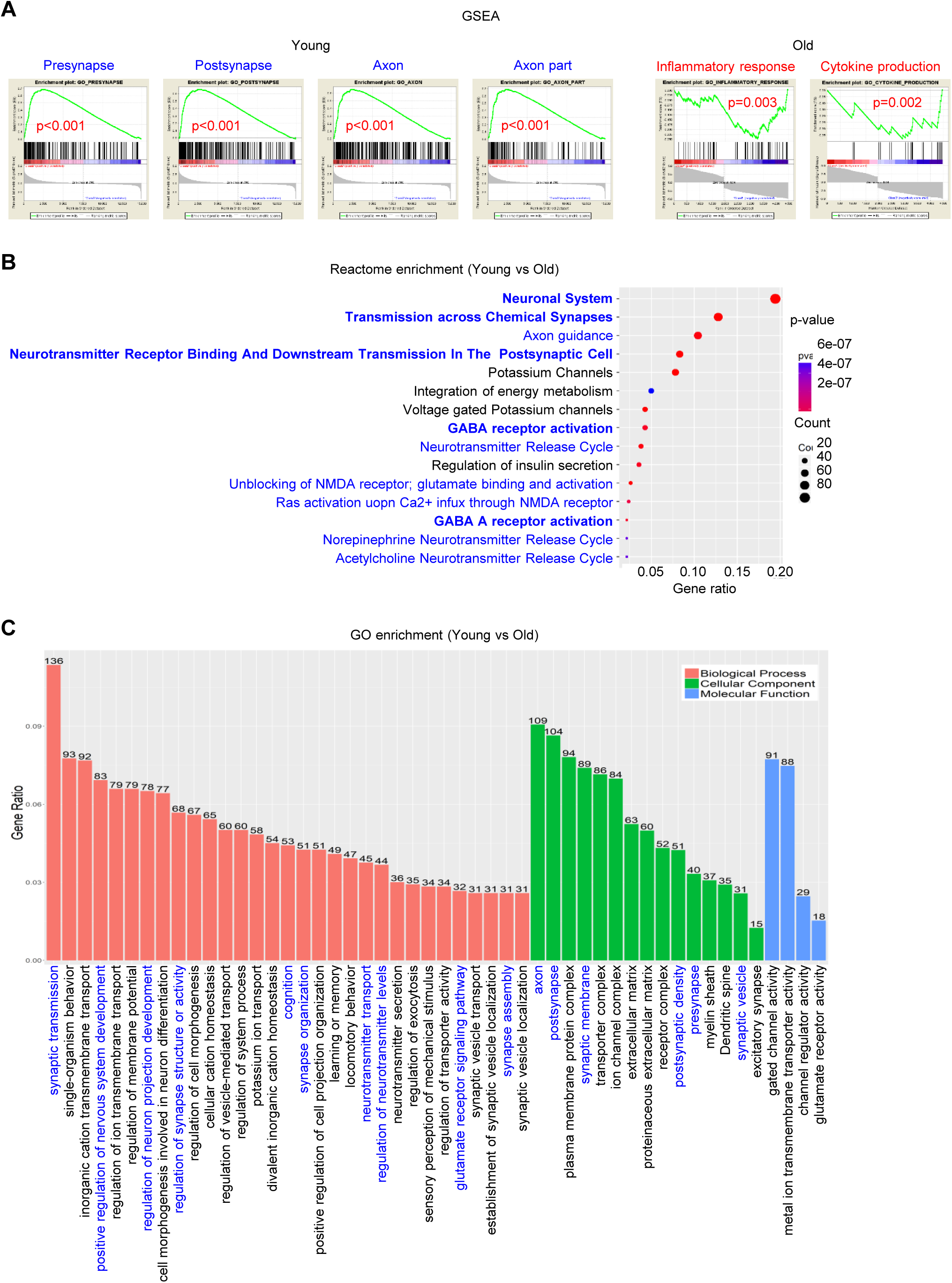
(**A**) Upregulated gene sets in addition to those in Figure 2E in young and old glioblastoma patients. Gene expression profile data were obtained from the TCGA database. (**B-C**) Reactome enrichment analysis (**B**) and Gene ontology (GO) pathway analysis (**C**) of RNA-seq data for comparison between young and old mouse tumors.

**Figure S7 (complements Figure 3).**
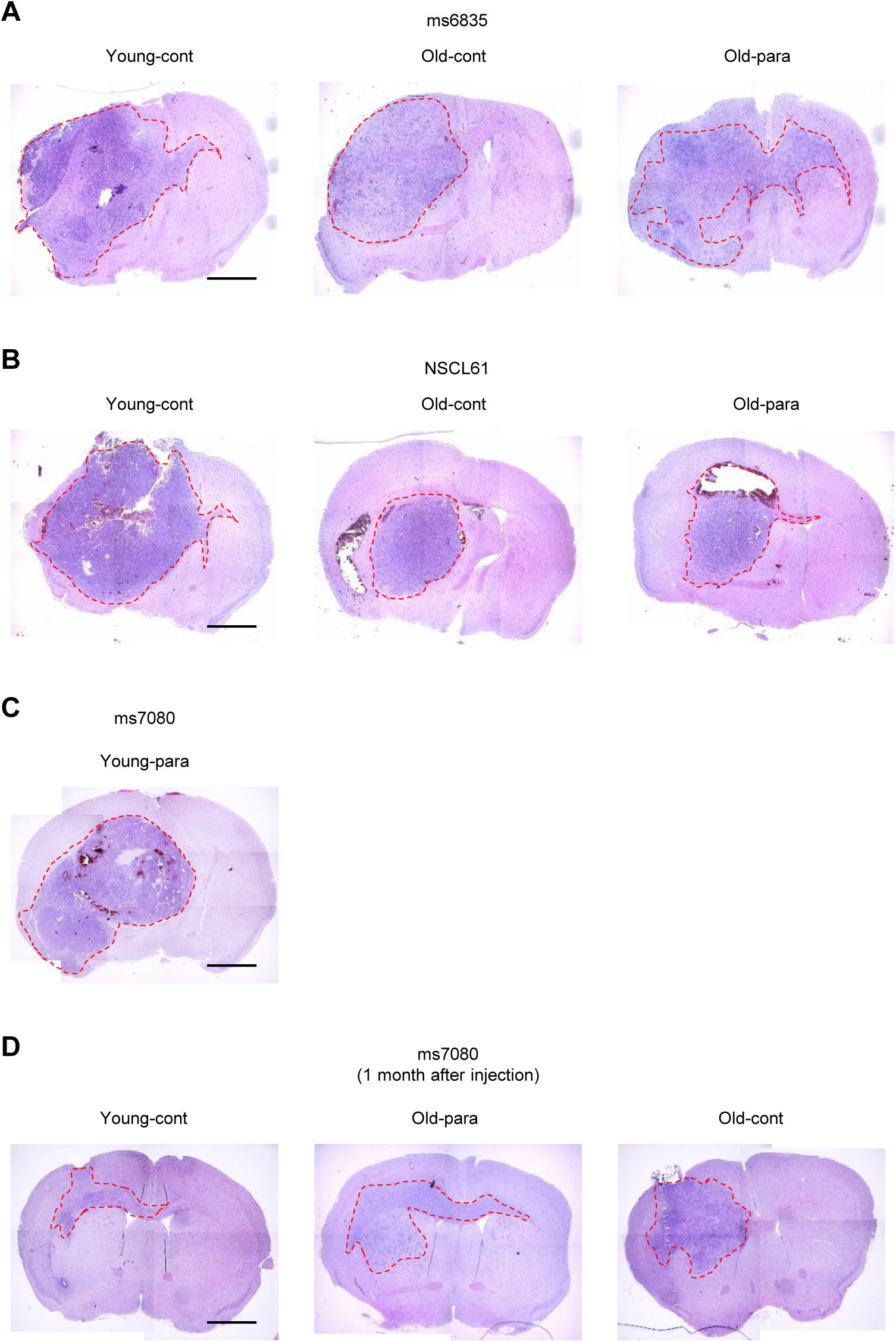
(**A-B**) H&E staining of Young-cont (top), Old-cont (middle) and Old-para (bottom) mouse brains following intracranial injection of ms6835 (A) and NSCL61 (B) cells. Scale bars represent 2 mm. (**C**) H&E staining of Young-para mouse brains following intracranial injection of ms7080 cells. Scale bars represent 2 mm. (**D**) H&E staining of young-cont (top left), old-cont (top right), old-cont (bottom left), and old-young (bottom right) mice brains 1 month after intracranial injection of ms7080 cells. Scale bar represents 2 mm.

**Figure S8 (complements Figure 3).**
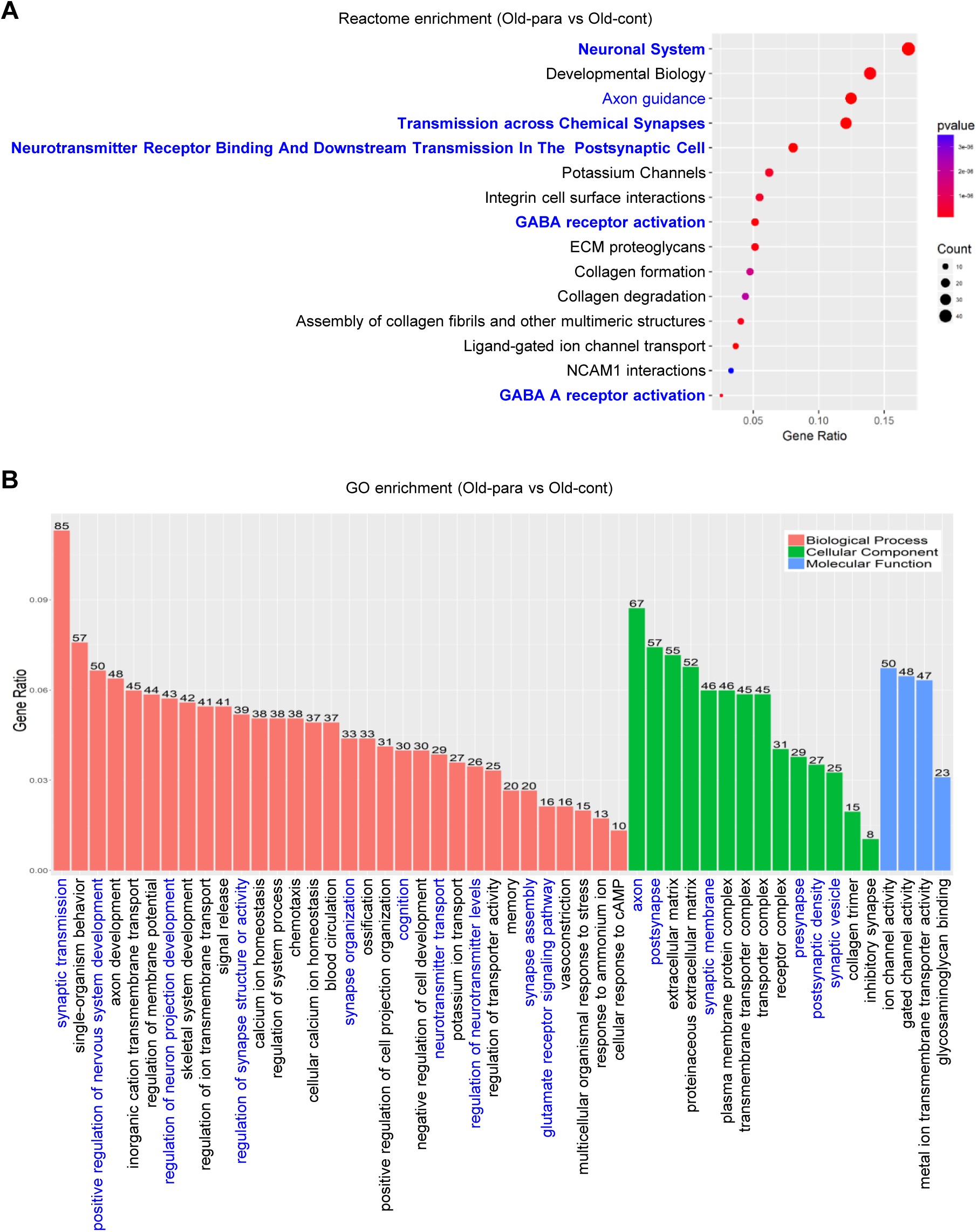
(**A-B**) Reactome enrichment analysis (**A**) and GO pathway analysis (**B**) of RNA-seq data for comparison between Old-para and Old-cont tumors.

**Figure S9 (complements Figure 4).**
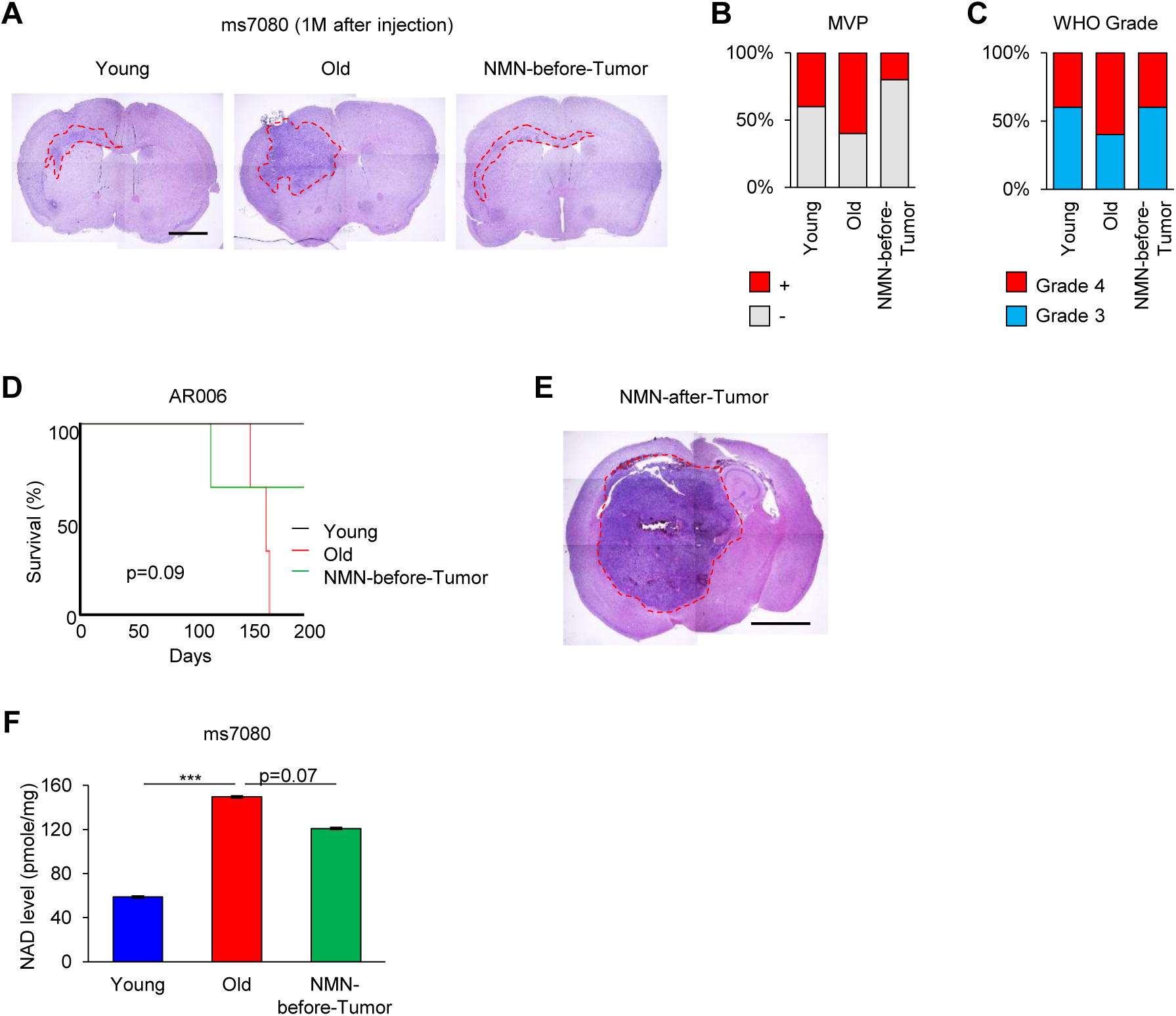
(**A**) Representative H&E staining of young, old, and NMN-before-tumor mice at one month after intracranial injection of ms7080 cells. Scale bar represents 2 mm. (**B**) Ratio of tumor tissues with microvascular proliferation (MVP). Data are the means ± SD (n=5 each). (**C**) WHO grading of tumor tissues in young, old, and NMN-before-tumor mice at one month after intracranial injection of ms7080 cells (n=5 for each). (**D**) Kaplan-Meier curve comparing survival of young, old, and NMN-before-tumor mice following intracranial injection of AR006 cells (n=3 for each). (**E**) H&E staining of ms7080-injected mouse brains together with post-injection NMN treatment (NMN-after-Tumor). Scale bar represents 2 mm. (**F**) Quantitative analysis of NAD^+^ level by LC/MS in ms7080 tumors in young, old, and NMN-before-tumor mice (n=5 each). \*\*\**p*<0.001.

**Figure S10 (complements Figure 4).**
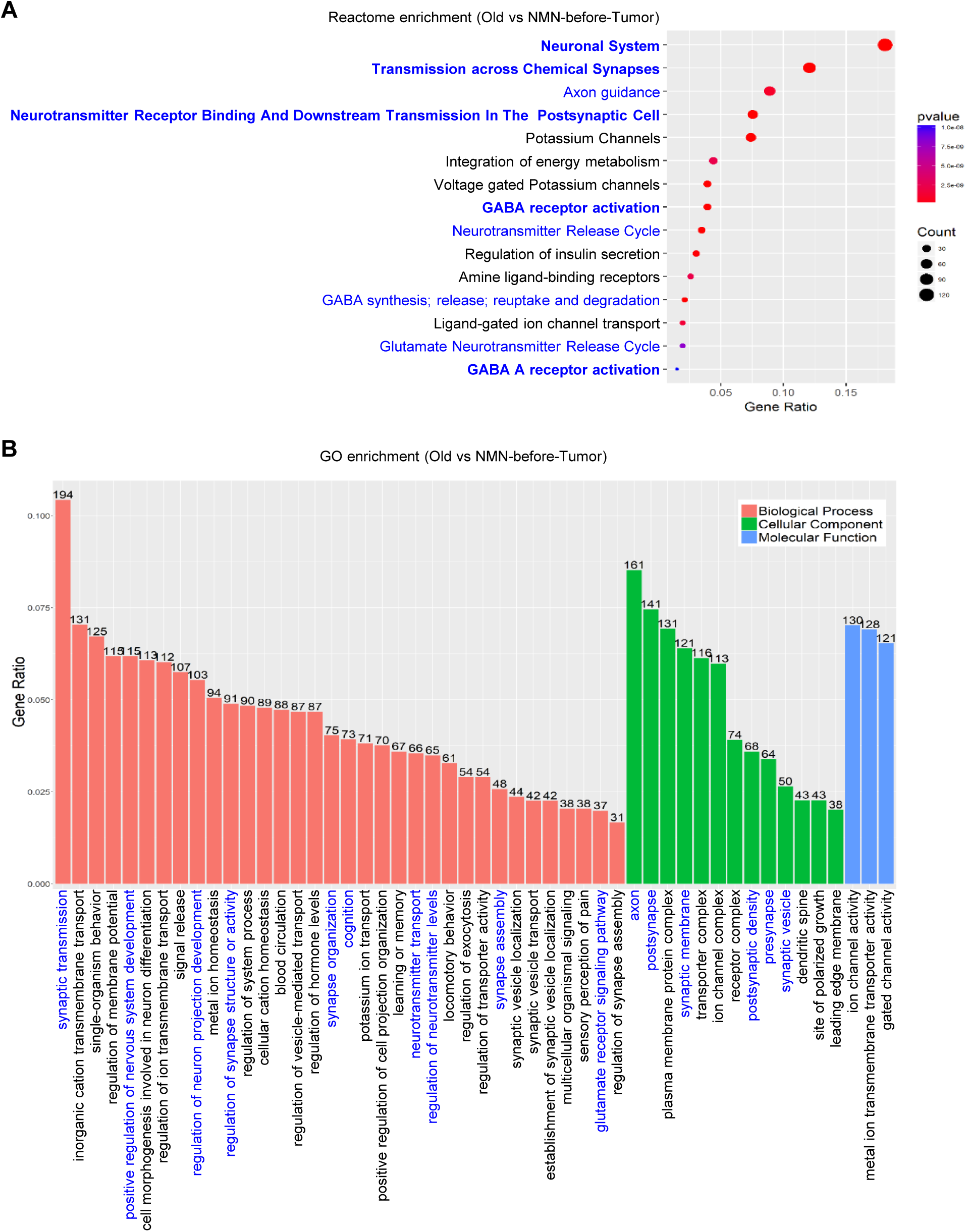
(**A-B**) Reactome enrichment analysis (**A**) and GO pathway analysis (**B**) of RNA-seq data for comparison between NMN-before-tumor and old tumors.

**Figure S11 (complements Figure 4).**
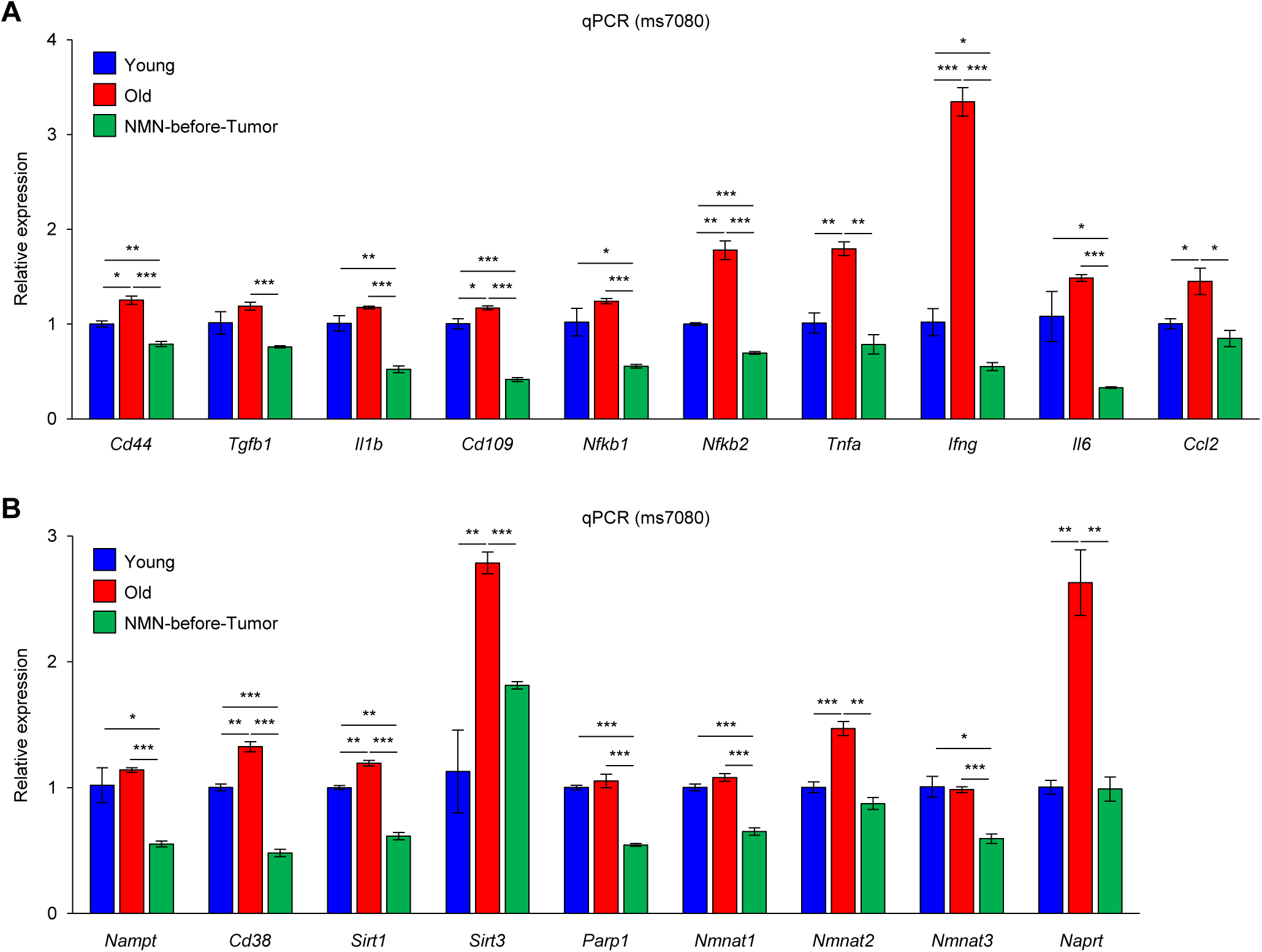
RT-qPCR analysis of mRNAs encoding neuroinflammation factors (**A**) and NAD+ pathway-related factors. (**B**) Gene expression programs in cortical tissues of young, old, and NMN-before-tumor mice. Data are the means ± SD (n=3 each).

**Figure S12 (complements Figure 4).**
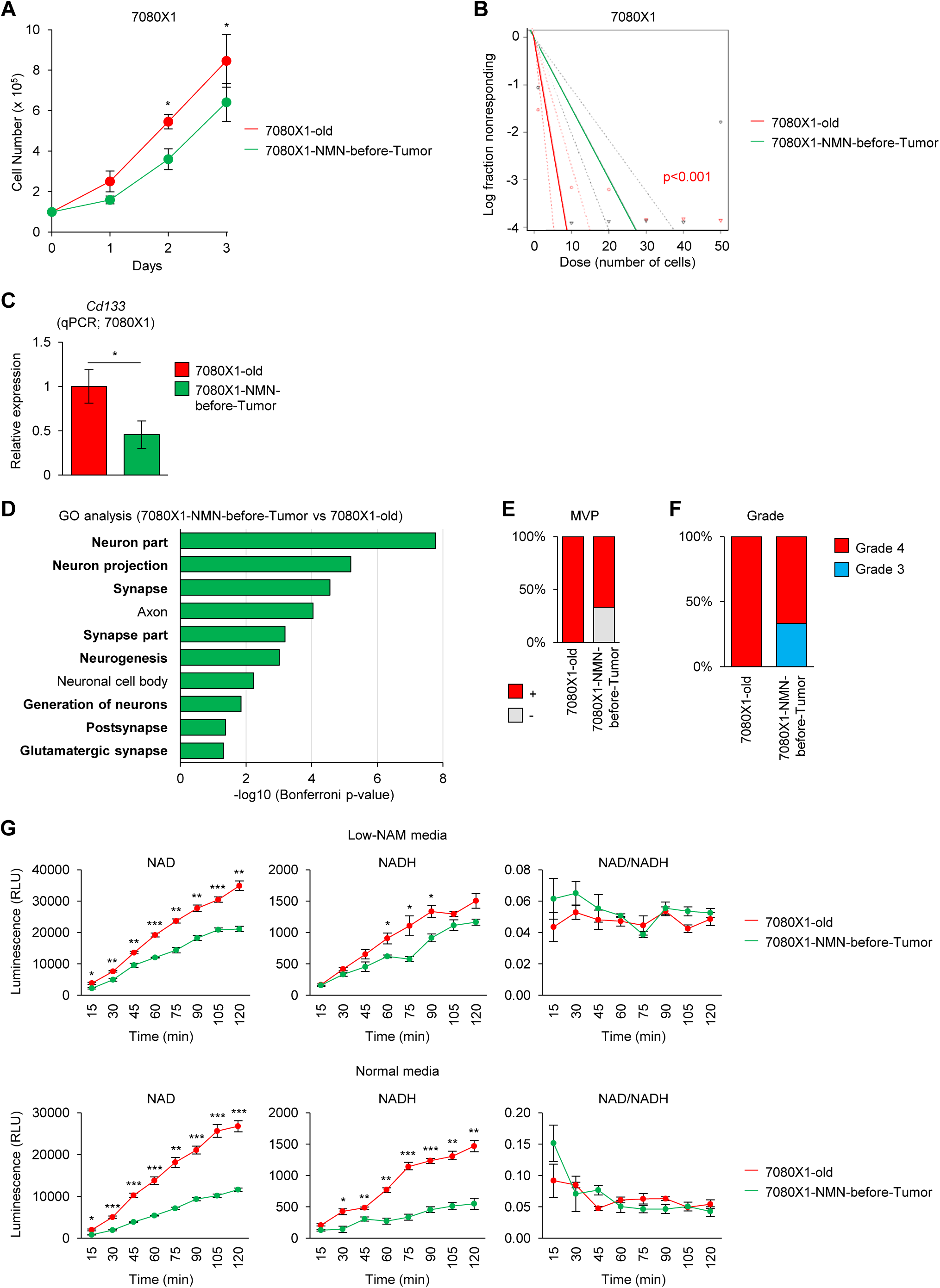
(**A**) Proliferation in culture of 7080X1-old and -NMN-before-tumor cells. Data are the means ± SD (n=3 each). \**p*<0.05. (**B**) Log-dose response curves of sphere formation assays of 7080X1-young, -old, and -NMN-before-tumor cells. Data are the means ± SD (n=5 each). (**C**) RT-qPCR analysis of the expression levels of *CD133* mRNA in 7080X1-old and -NMN-before-tumor cells. Data are the means ± SD (n=3). \**p*<0.05. (**D**) GO analysis of RNA-seq data showing the list of activated biological processes in 7080X1-NMN-before-tumors compared to 7080X1-old tumors. (**E**) Ratio of 7080X1 tumor tissues with microvascular proliferation (MVP). Data are the means ± SD (n=5 each). (**F**) WHO grading of 7080X1 tumor tissues at one month after injection (n=5 for each). (**G**) Quantitation of the levels of NAD^+^, NADH and NAD^+^/NADH in 7080X1-old and -NMN-before-tumor cells cultured in low-nicotinamide (NAM) media (top) and normal media (bottom), using luminescence [relative light units (RLUs)]. Data are the means ± SD (n=3 each).

**Figure S13 (complements Figure 5).**
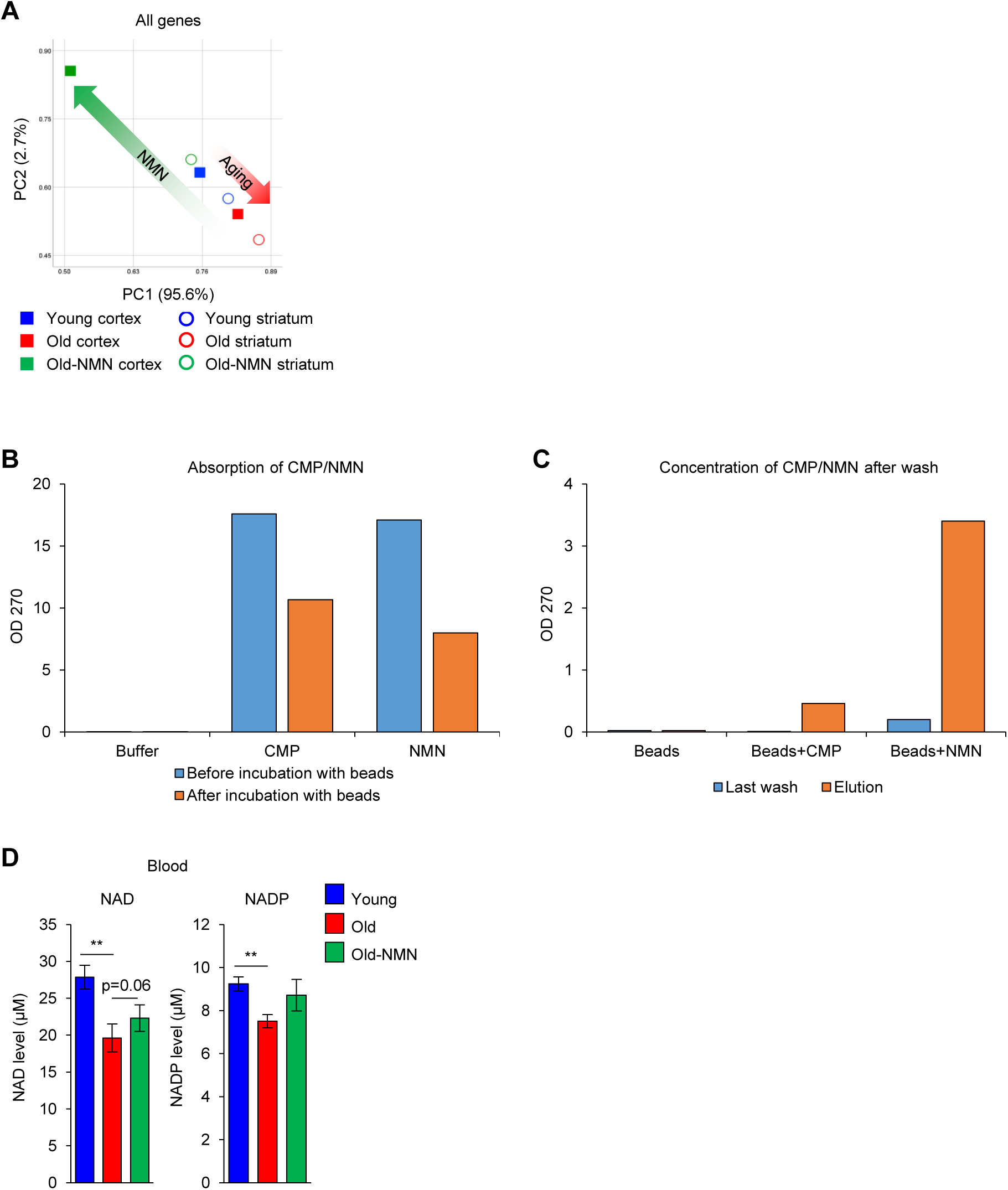
(**A**) PCA of RNA-seq analysis of young, old, and old-NMN cortical and striatum tissues. (**B**) CMP and NMN in solution after incubation with specific beads. (**C**) CMP and NMN attached to the beads after wash and elution. (**D**) Quantitative analysis of NAD^+^ and NADP levels in young, old, and old-NMN mouse blood samples. Data are the means ± SD (n=10 each). \*\**p*<0.01.

**Figure S14 (complements Figure 6).**
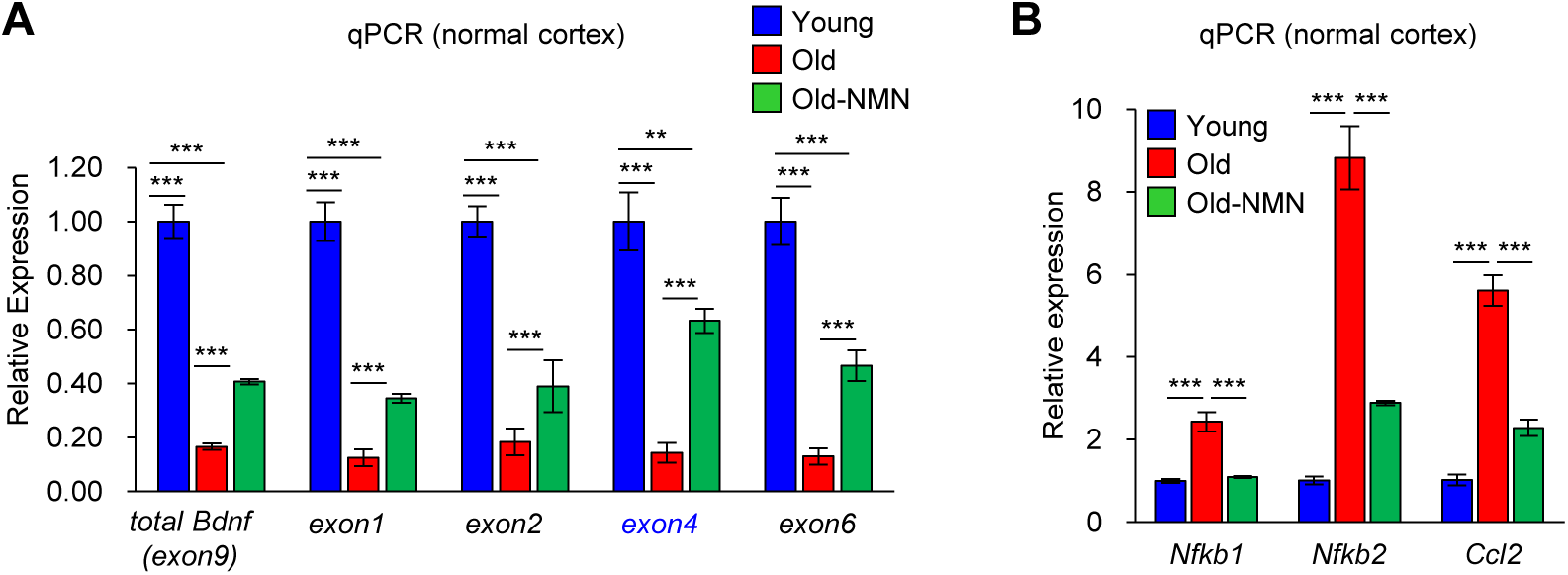
RT-qPCR analysis of *BDNF* mRNA as well as exons 1, 2, 4, and 6 of *BDNF* mRNA (**A**) and *Nfkb1*, *Nfkb2*, and *Ccl2* mRNAs, encoding neuroinflammatory markers (**B**), in young, old, and old-NMN cortices. Data are the means ± SD (n=6 each). \*\**p*<0.01, \*\*\**p*<0.001.

**Figure S15 (complements Figure 6).**
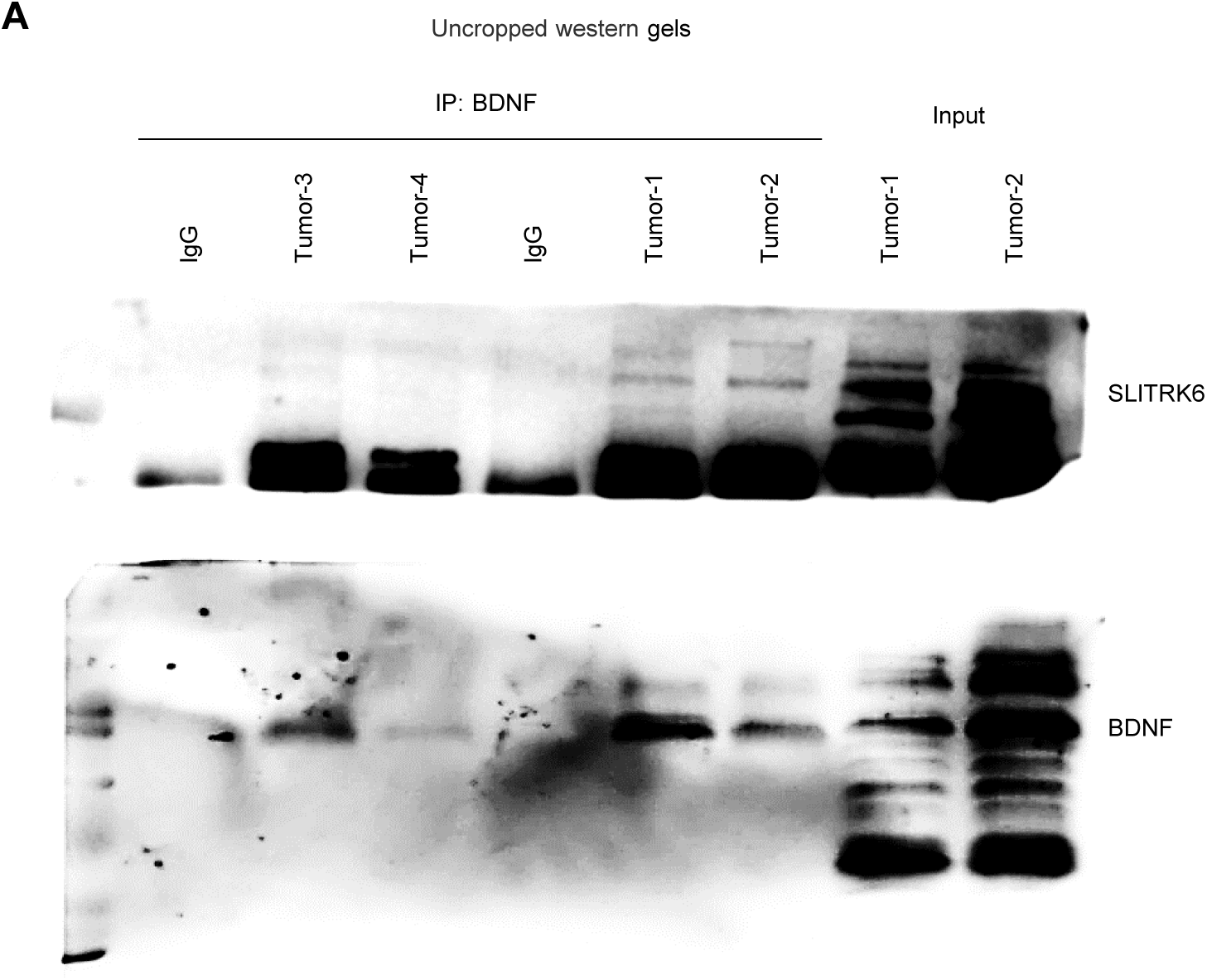
(**A**) Uncropped western gels for SLITRK6 (top) and BDNF (bottom).

